# Fast, long-range intercellular signal propagation through growth assisted positive feedback

**DOI:** 10.1101/2024.12.04.626899

**Authors:** Meidi Wang, Louis González, Soutick Saha, Krešimir Josić, Andrew Mugler, Matthew R. Bennett

**Affiliations:** PhD Program in Systems, Synthetic, and Physical Biology, Rice University, Houston, TX, USA; Department of Physics and Astronomy, University of Pittsburgh, Pittsburgh, PA, USA; Department of Physics and Astronomy, Purdue University, West Lafayette, IN, USA; Department of Biosciences, Rice University, Houston, TX, USA; Department of Mathematics, University of Houston, Houston, TX, USA; Department of Biology and Biochemistry, University of Houston, Houston, TX, USA; Rice Synthetic Biology Institute, Rice University, Houston, TX, USA; Department of Bioengineering, Rice University, Houston, TX, USA

**Keywords:** Genetic circuit engineering, intercellular signaling, mathematical modeling, trigger wave, synthetic biology

## Abstract

Intercellular signaling in bacteria is often mediated by small molecules secreted by cells. These small molecules disperse via diffusion which limits the speed and spatial extent of information transfer in spatially extended systems. Theory shows that a secondary signal and feedback circuits can speed up the flow of information and allow it to travel further. Here, we construct and test several synthetic circuits in *Escherichia coli* to determine to what extent a secondary signal and feedback can improve signal propagation in bacterial systems. We find that positive feedback-regulated secondary signals propagate further and faster than diffusion-limited signals. Additionally, the speed at which the signal propagates can accelerate in time, provided the density of the cells within the system increases. These findings provide the foundation for creating fast, long-range signal propagation circuits in spatially extended bacterial systems.

## INTRODUCTION

Intercellular signaling plays an essential role in both unicellular and multicellular organisms. Intercellular communication in multicellular organisms is critical for coordinating cell differentiation during development^1,2^. In unicellular organisms, such as bacteria or yeast, intercellular signaling can help colonies synchronize their behavior to improve survival in challenging and changing environments^3–6^. In synthetically engineered multicellular systems, intercellular communication plays a central role in the emergence of programmed spatiotemporal patterns within the population^7–10^.

In bacteria, intercellular signals are mediated by a variety of molecules^11^, such as acylhomoserine lactones^12^, quinolones^13^, and nitric oxide^14^ that, once secreted or transported out of a cell, diffuse through the extracellular space. Although diffusion is fast and reliable across distances on the order of the size of a single cell, it becomes inefficient for propagating information at millimeter or centimeter scales^15,16^. At large distances diffusion is relatively slow, and the strength of the signal (*i*.*e*. its local concentration) rapidly decreases as a function of distance from the source(Fig. 1A).

**Figure 1:**
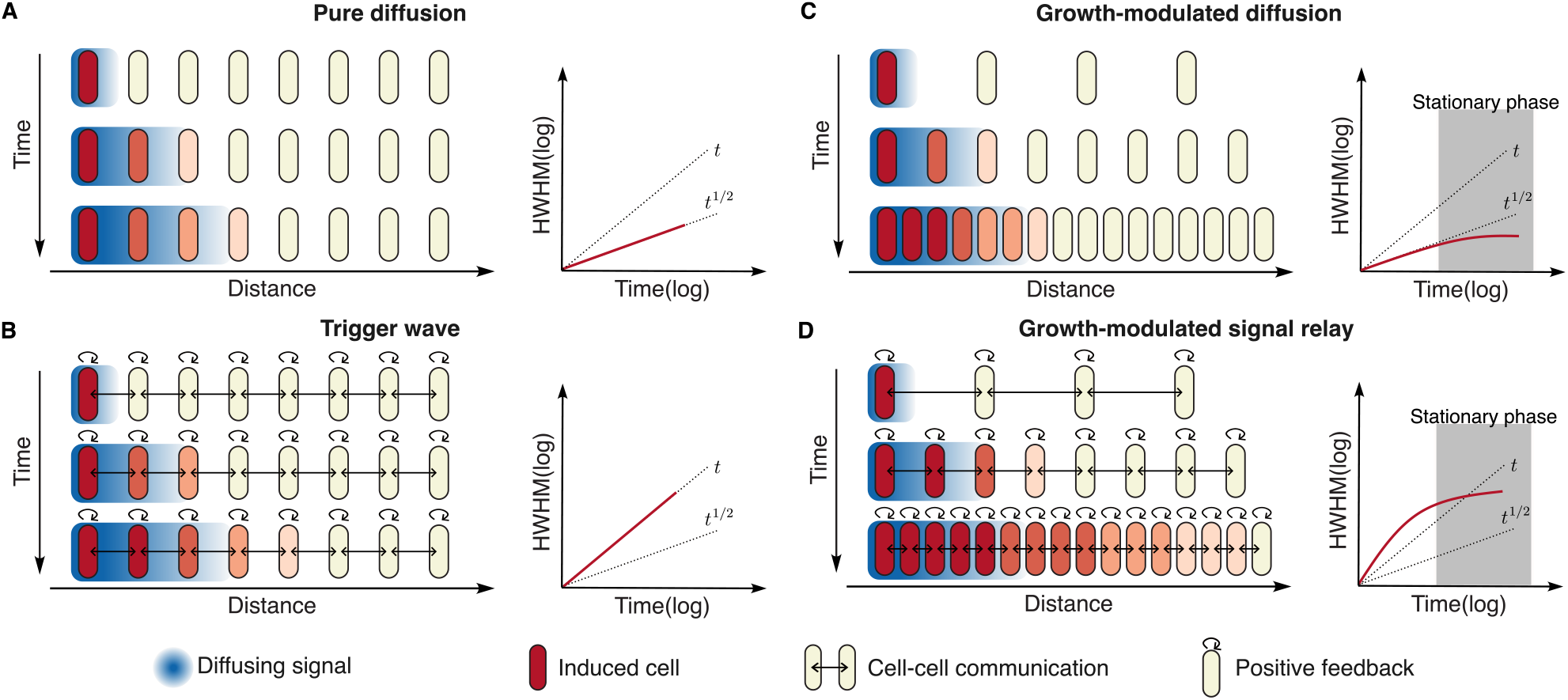
Conceptual illustration of how bacteria respond to a local signal. (**A**) Pure diffusion: The signal diffuses as a gradient (blue) and activates a cellular response (red). Half-Width Half-Maximum (HWHM), the distance from the maximum signal response to the point where the signal response falls to half of its maximum value, represents the extent of signal propagation. Cells show a diminishing response over time and distance. (**B**) Trigger wave: Incorporates both spatial coupling and feedback loops, as indicated by arrows between and within cells. Each induced cell (red) stimulates neighboring cells, thereby maintaining a constant signal propagation speed. (**C**) Growth-modulated diffusion: Similar to pure diffusion but includes the impact of cell growth dynamics on signal propagation. The signal propagates similarly to pure diffusion and stops as cells approach the stationary phase. (**D**) Growth-modulated signal relay: A combination of cell growth and intercellular positive feedback. Active cell growth accelerates signal propagation before it slows down when cells enter stationary phase.

Cells can overcome the limitations of diffusion and coordinate behavior over large distances by combining intercellular signaling mechanisms with positive feedback to create spatial trigger waves^17^. Such waves maintain their amplitude and speed during signal propagation^18,19^ (Fig. 1B). While spatial trigger waves are typically associated with metazoans, similar mechanisms may play a role in signal propagation within bacterial communities^20,21^. A self-amplifying potassium wave in *Bacillus subtilis* represents an electrical signaling mechanism that propagates through biofilms to coordinate resources^22^. Additionally, bacterial quorum sensing (QS) systems, which are well-established for sensing cell density^23^, are also suggested to function as chemical signaling waves that mediate intercellular communication in spatially heterogeneous environments ^21,24–26 27^. Studies have suggested that autoinducer diffusion, coupled with positive feedback-regulated autoinducer synthesis, facilitates signal propagation in a traveling wave fashion^20,28,29^. Indeed, experiments have shown that distant cells are activated when local QS molecules are supplied exogenously^21,30–32^.

However, in native QS systems, auto-inducer production ties to population density, so it emerges only after a quorum is reached. Long-distance communication through the QS system in bacteria is particularly complex because signal propagation through gene expression is inherently coupled with bacterial growth and division ^21,30,33,34^. The exponential growth of bacteria may act as an amplifier of signal propagation as cell density increases. On the other hand, once cells reach stationary phase, they exhibit sharply reduced growth and gene expression, damping both the production and amplification of intercellular signals (Fig. 1C,D). Autoinducers have been shown to accelerate the diffusing signal front^21,30^, but only when added exogenously to the bacteria performing quorum sensing. As population size increases, the self-activation of autoinducers complicates the ability to distinguish whether true signal propagation is occurring or if the system is merely being triggered by reaching a population threshold^21^. This leaves it unclear whether an auto-inducer traveling wave can be engineered to propagate signals in response to a local stimulus, independent of a global population threshold. Thus, a new experimental approach and an accompanying theoretical framework are needed to understand the complex interplay between cellular population growth, gene expression, and signal transmission.

To investigate the role of positive feedback in long-range intercellular signaling in growing bacterial communities, we constructed a synthetic signal relay system in *E. coli* that uses positive feedback to amplify and relay signals as they propagate through a spatially extended community. This bottom-up approach allowed us to systematically probe each component of the signaling system to determine how they affect a community’s response to a spatially localized input signal. Importantly, the positive feedback in our engineered bacteria does not lead to self-activation at high cell densities. This is in contrast to QS systems found in nature that, due to leaky expression of the AHL synthase, auto-activate at large cell densities^26^. We could thus decouple diffusion and feedback-mediated signal propagation from population mediated signal activation.

We found that *E. coli* with a positive feedback-coupled intercellular signaling system exhibited super-diffusive signal propagation, and that such propagation was not possible when either the feedback or the intercellular signaling was removed. Furthermore, we found that exponential growth of the cells themselves enhanced propagation beyond the constant-speed ballistic regime, where distance increases linearly with time, and that the propagation rate could be tuned by altering the duration of the exponential growth phase (Fig. 1D). A mathematical model explained why both feedback and intercellular signaling are required for robust super-diffusive propagation, and it revealed how exponential cell growth can lead to hyperballistic signal propagation (*i*.*e*. signals that accelerate with time). We thus demonstrate that a bacterial colony can overcome the short length scales of diffusive communication using positive feedback and intercellular signaling, and can surpass even ballistic propagation by coupling signaling and population growth.

## RESULTS

### Growth-modulated signal relay

To investigate the roles of growth and feedback on intercellular signaling, we engineered a strain of *E. coli* (IS+PF+) that responds to an externally applied input signal (erythromycin) and uses a secondary signal (C4 homoserine lactone, C4-HSL) and feedback control to spatially propagate the response throughout the population (Fig. 2A). The addition of erythromycin de-represses the P_mph_ promoter which drives expression of *rhlI* ^35^, a gene encoding the acyl-homoserine lactone synthase, RhlI. When present, RhlI produces C4-HSL, a small molecule that freely diffuses in and out of cells and binds to the transcriptional activator RhlR to up-regulate transcription from P_rhl_ promoters^36^. To incorporate positive feedback without causing self-activation, we utilized a leak-dampener module previously developed by Ho et al.^37^. This module, driven by P_rhl_, includes a second copy of the *rhlI* gene, denoted as *rhlI*.*2TAG*, with the first two leucine codons mutated to amber stop codons (TAG), and the leucine suppressor tRNA *supP*. In the absence of C4-HSL, the translation of *rhlI*.*2TAG* is significantly reduced by the stop codons. In the presence of C4-HSL, *supP* enables translation of *rhlI*.*2TAG*, which produces more C4-HSL. Both RhlI and RhlI.2TAG contain C-terminal ssrA degradation tags that target the ClpXP protease^38^. An additional copy of P_rhl_ drives expression of the gene encoding superfolder yellow fluorescent protein (sfYFP), which we use to monitor the response of the cells. *mphR* and *eryR* are in the same expression cassette controlled by the constitutive promoter P_lacIq_ ^35^. The plasmids containing this circuit were transformed into *E. coli* strain CY027 with *rhlR* encoded in the genome^8^.

**Figure 2:**
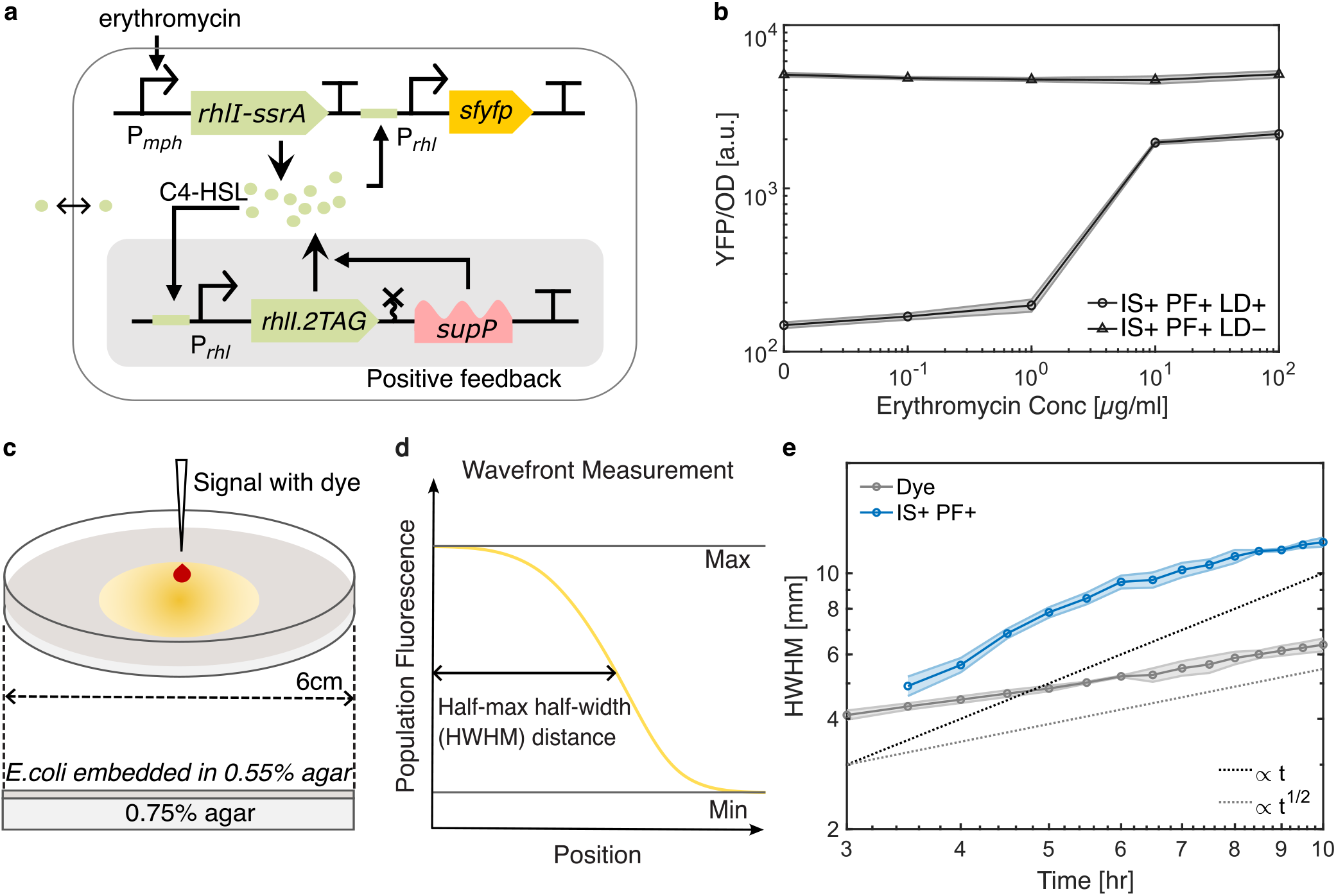
Signal relay strain propagates signals super-diffusively. (**A**) Circuit diagram of the signal relay strain (IS+PF+). Erythromycin increases C4-HSL production via the RhlI synthase. C4-HSL induces the expression of *sfyfp, rhlI*.*2TAG*, and the transcription of *supP*. The *rhlI*.*2TAG* has two leucine codons replaced by amber stop codon TAG. Increased transcription of *supP* allows for more *rhlI* expression by mediating correct translation of leucin codons. (**B**) Fluorescence intensity of the IS+PF+ strain in liquid culture, comparing conditions with and without the leak dampener (LD) incorporated in the positive feedback module. The leak dampener reduces auto-activation of the IS+PF+ strain. (**C**) Schematic of experiment design illustrating cell culture in solid medium and application of exogenous signal. (**D**) Wavefront quantified using the half-width at half-maximum (HWHM) of the fluorescence profile. Maximum and minimum fluorescence levels are identified at each time point. Wavefront distance is measured at half of the fluorescence range (max – min). (**E**) Measured wavefront dynamics (half width half maximum HWHM) of the growth-modulated IS+PF+ strain and the dye. Shaded region corresponds to one standard deviation of three technical replicates. Both axes are set to a logarithmic scale.

We next confirmed that the IS+PF+ strain can respond to an external erythromycin signal and report with *sfyfp* expression. We tested the strain in liquid culture across a range of erythromycin concentrations (Fig. S1A). As expected, higher concentrations of erythromycin resulted in increased sfYFP expression (Fig. S1A). The addition of erythromycin induced fluores-cence expression without affecting the growth rate. To ensure that positive feedback does not cause self-activation in the absence of erythromycin, we grew the IS+PF+ strain in liquid and solid media and compared its fluorescence with two control strains: one with intercellular signaling but without positive feedback (IS+PF–), and another with intercellular signaling but lacking the leak dampener systems (IS+PF+LD–). Similar to the IS+PF+ strain, the IS+PF– strain (Fig. S2A) contains the P_mph_ promoter driving expression of *rhlI* and the P_rhl_ promoter driving expression of *sfyfp*. However, the IS+PF– strain does not contain the *rhl*.*2TAG* and hence there is no positive feedback on *rhl* expression. Note that the second copy of the P_rhl_ promoter still drives transcription of *supP*. When grown in LB broth and soft LB agar, the fluorescence levels of both IS+PF+ strain and IS+PF– strain were comparable, suggesting no self-activation was introduced by the leak dampened positive feedback. In contrast, the IS+PF+LD– strain exhibited high levels of fluorescence without erythromycin, indicating self-activation (Fig. 2B, Fig. S1B). Together, these results confirm that the IS+PF+ strain is tunable with erythromycin and does not exhibit self-activation due to the positive feedback loop.

To determine if the IS+PF+ strain can propagate erythromycin signals faster than diffusion, we embedded exponentially-growing cells into 0.55% soft-agar and applied erythromycin with a red fluorescent dye (Sulforhodamine 101) to the center of the plate (Fig. 2C). We recorded the fluorescence on the plate every 30 minutes for 10 hours after inoculating the cells (see Methods) and quantified the wavefront movement by the half-width half-maximum (HWHM) of the signal (Fig. 2D). Figure 2E showed that the wavefront of sfYFP in the IS+PF+ strain outpaced the wavefront of the dye. As expected for a diffusive molecule, the diffusion of the dye led to an increase in wavefront distance roughly proportional to the square root of the elapsed time. There is a small deviation from the theoretical scale, likely due to the non-uniform loss of fluorescence dye to the bottom layer of the agar. In contrast, the wavefront of sfYFP traveled approximately linearly with time before eventually slowing down.

We hypothesized that the slowdown observed at later times occurred due to cells entering stationary phase, which is characterized by nutrient or space depletion leading to, among other effects, an equilibrium between cell division and cell death and changes in gene expression^39,40^. During stationary phase, *E. coli* activates gene expression related to maintenance of resource and energy for survival and downregulates other metabolic activities, resulting in reduced production of C4-HSL and sfYFP. We verified the growth phase of cells on agar plate by measuring the cell density change over the course of the experiment and confirmed that cell density saturated around five hours after we seeded them in agar, suggesting a growth phase shift (Fig. S3).

### Intercellular signaling and positive feedback in signal relay

We next asked whether both intercellular signaling and positive feedback are necessary for signals to propagate faster than diffusion. To answer this question, we created two new strains – one incorporating positive feedback, but lacking intercellular signaling (IS–PF+), and one lacking both intercellular signaling and positive feedback (IS–PF–). For the IS–PF+ strain (Fig. S2B), the P_mph_ promoter controls the expression of the gene encoding the transcriptional activator *araC* instead of *rhlI*. AraC, when bound to arabinose, activates its target promoter (P_BAD_)^41^ and, importantly, does not synthesize a diffusible signaling molecule like RhlI does. Abundant arabinose is provided in the system. There are two copies of P_BAD_, one driving expression of the reporter *sfyfp* and the other driving expression of a leak-dampened version of *araC, araC*.*2TAG*. Both AraC and AraC.2TAG contain C-terminal ssrA degradation tags that target the ClpXP protease^38^. The IS–PF– strain (Fig. S2C) contains neither intercellular signaling nor positive feedback. This strain contains the P_mph_ promoter driving expression of *araC* and two copies of P_BAD_, one driving *sfyfp* and the other *supP* (without *araC*.*2TAG*). We measured the response of the four circuits to various erythromycin concentrations in liquid culture (Fig. S1A). All strains showed similar sensitivity to erythromycin. The strains with positive feedback exhibited higher maximum fluorescence while maintaining low basal expression, suggesting low self-activation. The basal expression level of the IS–PF+ strain was also confirmed in soft LB agar. The fluorescence level of the IS–PF+ strain was similar to the fluorescence level of the IS–PF– strain when the leak dampener is incorporated (Fig. S1B).

After confirming that each strain functions properly in a homogeneous environment, we performed signal propagation assays as described above. Exponentially growing cells were embedded in soft agar and erythromycin was deposited at the center of the plate. Time-lapse images showed that the IS–PF– strain exhibited the lowest fluorescence, due to a weaker fluorescence expression cassette (Fig. 3A). The IS+PF– strain exhibited substantially higher fluorescence, although it showed similar levels of sfYFP expression to the IS–PF– strain in liquid culture (Fig. S1A). The difference in fluorescence between liquid and solid media was due to the fast dilution of C4-HSL in liquid media caused by the circular shaking motion of liquid media^42,43^. This faster dilution led to lower activation of the P_rhl_ promoter than the same cells in solid media. The IS– PF+ and IS+PF+ strains showed higher fluorescence level than the strains without the positive feedback, consistent with results in liquid culture (Fig. S1A).

**Figure 3:**
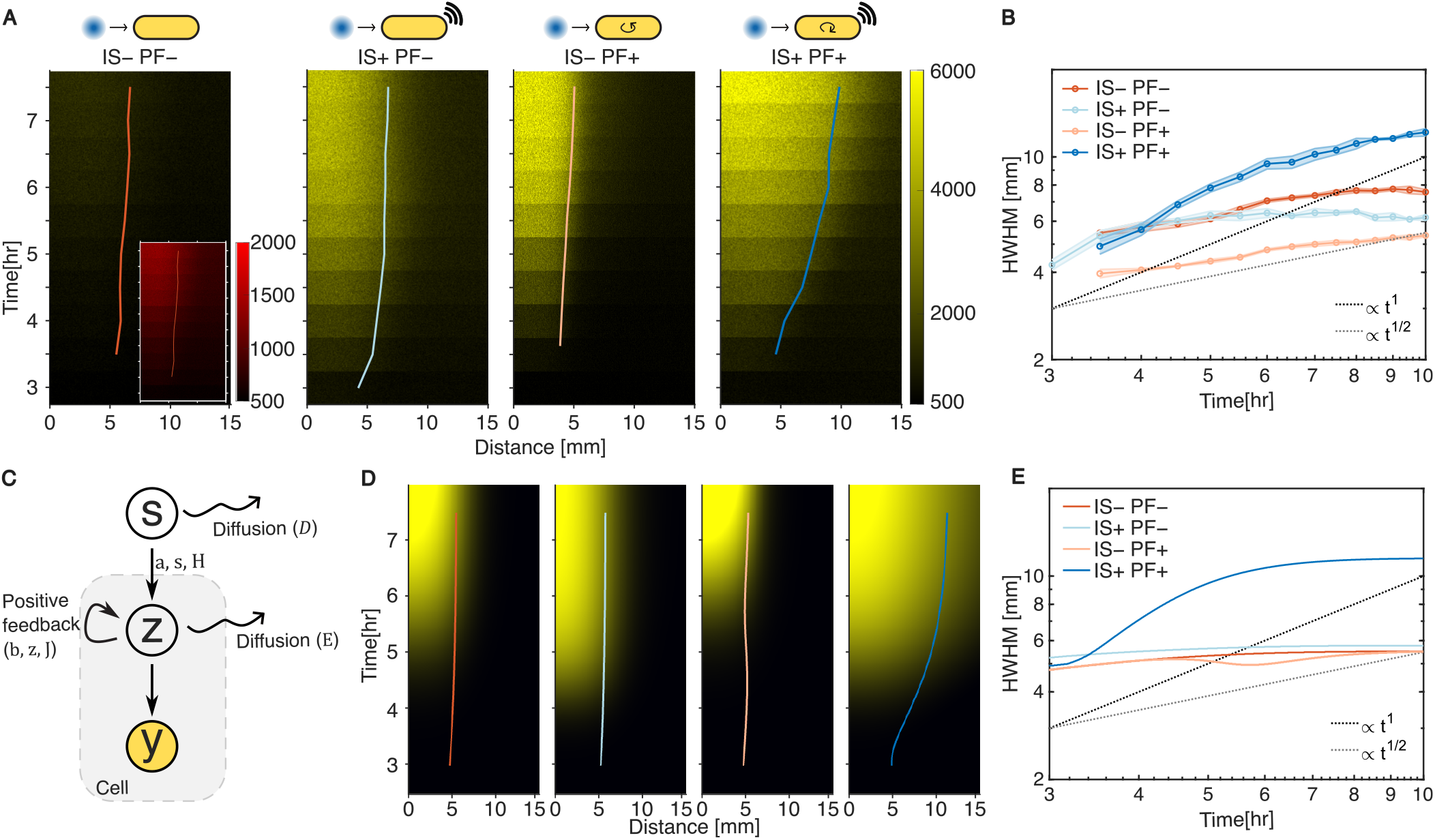
Signal propagation dynamics in strains with and without intercellular signaling and positive feedback. (**A**) Time-lapse montage of sfYFP in the IS–PF–, IS+PF–, IS–PF+, and IS+PF+ strains. Fluorescence intensities are normalized to the same maximum and minimum values. The inset shows the IS–PF– strain with fluorescence intensities normalized to lower maximum values, represented by the red color bar on the right. (**B**) Measured wavefront dynamics (HWHM) in four strains. Shaded region corresponds to one standard deviation of three technical replicates. Both axes are set to a logarithmic scale. (**C**) Basic logic of the mathematical model. Diffusing erythromycin (*s*) induces the production of AraC or C4-HSL, denoted as *z*. Depending on the circuit design, *z* may have positive feedback and diffuse. *z* activates the production of sfYFP (*y*), and both *z* and *y* production are modulated by cell growth. (**D**) Simulated sfYFP profiles of each strain with the HWHM wavefront distance overlaid. The simulation parameters are listed in Table S1. Simulated fluorescence profiles qualitatively match the experimental results. (**E**) Simulated wavefront dynamics. The model and experiment data qualitatively agree. Both demonstrate the super-diffusive signal propagation in the IS+PF+ strain and diffusive propaga- tion in the IS–PF– strain, IS+PF– strain and IS–PF+ strain.

Signal propagation that relies solely on erythromycin faces the same limitations as the dye: the distance increases as the square root of time. To determine whether both intercellular signaling and positive feedback are necessary for the observed super-diffusive signal propagation, we measured the wavefront movements in each of the four strains. The IS+PF+ strain displayed elevated fluorescence expression at a substantially larger distance from the source (Fig. 3A,B). In contrast, the wavefront distances for the IS–PF–, IS–PF+, and IS+PF– strains increased with the square root of time, consistent with diffusion limits. Although the IS–PF+ strain showed higher fluorescence than the IS–PF– strain due to positive feedback, its absolute wavefront distance was actually smaller. This can be explained by the increased fluorescence intensity at the diffusing source and by the fact that this amplified response did not spread to neighboring cells due to the lack of intercellular signaling. The IS+PF– cells responded to erythromycin with C4-HSL production, which activates *sfyfp* expression. Although C4-HSL diffusion smoothed the transition between activated and nonactivated cells, its limited diffusion restricted the overall propagation. Thus, only the IS+PF+ strain achieved super-diffusive signal propagation, effectively overcoming the erythromycin diffusion limitation.

### A mathematical model of signal propagation

To provide a mechanistic understanding of the experimental results and explain and predict the impact of changes in experimental conditions, we developed a mathematical model of the engineered system (Fig. 3C). We modeled the propagation of signaling molecules and the cellular responses using a system of partial differential equations (PDEs). Working in polar coordinates, and acknowledging the angular symmetry of the bacterial colony, we write the PDEs in terms of the radial coordinate, *r*, which measures the distance from the center of the population. These equations describe the concentration of the diffusing source, erythromycin (*s*), which induces the production of either AraC or C4-HSL (*z*), in turn, this signaling molecule produces the readout, sfYFP (*y*). The dynamics of these concentrations are described by

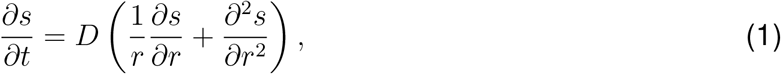

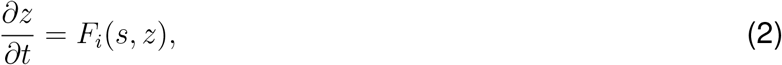

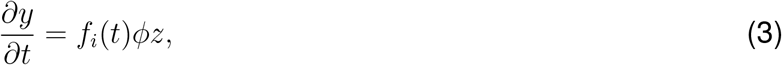

with the initial condition *s*(*r, t* = 0) = *s*_0_Θ(*r < R*_0_) and *z*(*r*, 0) = *y*(*r*, 0) = 0, where *s*_0_ is the initial concentration of erythromycin, *R*_0_ sets the radius of the initial droplet of erythromycin added to the medium, and Θ denotes the Heaviside step function. In equation (1), the right-hand side is the radial part of the 2-D Laplacian in polar coordinates with *D* the diffusion constant of erythromycin, and in equation (3) *φ* is an activation strength of *y* (sfYFP) by *z* (either AraC or RhlI). We use an absorbing boundary condition at r = 30 mm (Fig. 2B), but the specific choice of boundary condition here does not significantly influence the dynamics because the wave remains far from this boundary at all times (Fig. 3).

In equation (2), the function *F*_*i*_(*s, z*) captures the activation and feedback in each of the four strains. Specifically, in the IS–PF– case, 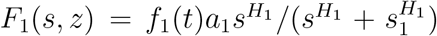 models the activation of AraC by erythromycin, with activation strength *a*_1_, Hill coefficient *H*_1_, and half-maximal concentration *s*_1_. In the IS–PF+ case, 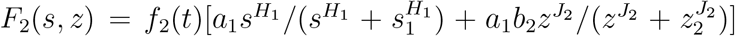 models both activation and feedback of AraC, with relative feedback strength *b*_2_, Hill coefficient *J*_2_, and half-maximal concentration *z*_2_. We assumed that the activation parameters *a*_1_, *H*_1_, and *s*_1_ are equal in the case of the IS–PF– and IS–PF+ strain because the activation components have the same molecular construction. In the IS+PF– case, 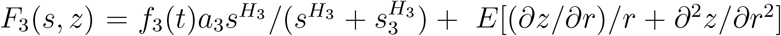 models the activation and diffusion of C4-HSL, which is synthesized by rhlI, with diffusion coefficient *E*. Finally, in the IS+PF+ case, 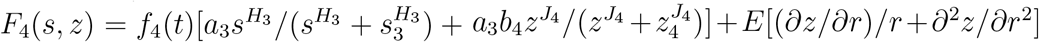 models the activation, feedback, and diffusion of C4-HSL. We again assumed that the activation parameters *a*_3_, *H*_3_, and *s*_3_ are equal for the IS+PF– and IS+PF+ strain because the induced production of C4-HSL by erythromycin is identical. Both *s* and *z* are treated as freely diffusing species. This simplification is justified because membrane equilibration of short-chain AHLs such as C4-HSL occurs in less than 30 s^44,45^. At each location, the transcriptional response is effectively in quasi-steady state with the local extracellular signal level. The net effect of intracellular binding events is absorbed into the Hill activation terms. The Hill coefficients and half activation term were calibrated directly to experimental data or analyzed systematically, providing a functional description of signal and promoter interactions. Our model is phenomenological, ignoring explicit delays due to multistep reactions, but we note that such delays can have important effects when coupled to feedback^46^, particularly on wave propagation^47,48^.

In equations (2) and (3), protein production is modulated by an envelope function, *f*_*i*_(*t*). This function reflects the fact that the rate of protein production depends on the rate of cell growth ^33^. Specifically, we assume that the growth envelope function is given by 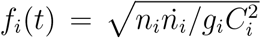 (Methods) where *n*_*i*_(*t*) is the cell number, *g*_*i*_ is the growth rate of the cells, and *C*_*i*_ is the carrying capacity for a given circuit *i*. The envelope function restricts protein production to the period of significant cell growth, after the lag phase and before stationary phase (Fig. S4). We assume that the number of cells per unit volume in the population grows logistically with a lag phase^49^, and has the form

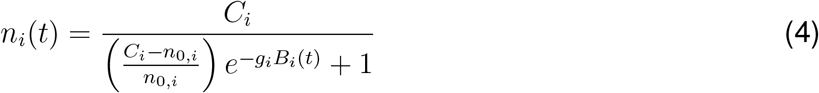

where *n*_0,*i*_ is the initial cell number and

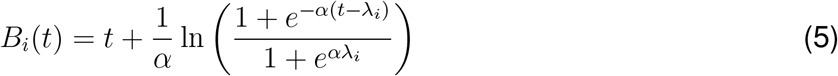

quantifies the cell growth from an initial lag phase with lag time *λ*_*i*_.

We obtained the parameters in equations (1-5) directly from our experimental measurements (*R*_0_, *s*_0_, *g*_*i*_, *n*_0,*i*_, *C*_*i*_, *λ*_*i*_, *a*_*i*_, *H*_*i*_, and *s*_*i*_) or from the literature (*D* and *E*), as described in the Methods. Where this was not possible (*J*_*i*_, *z*_*i*_, and *b*_*i*_), we explored the effects of these remaining parameters on the propagation dynamics and calibrated the parameters to the experimental dynamics (Methods). We also explored a version of the model with degradation of the intermediate (AraC is actively degraded whereas C4-HSL is not) and found that such degradation does not significantly affect the results (Methods). All parameter values are listed in Table S1.

We found that the mathematical model produces dynamic concentration profiles of sfYFP for all four strains (Fig. 3D) that agree qualitatively with those measured in the experiments (Fig. 3A). In particular, the model generates a traveling wave that remains high once activated and recapitulates the faster and longer signal propagation in the IS+PF+ compared to the other three strains. Figure 3E shows the HWHM propagation curves from the model, and we see excellent agreement with experimentally obtained propagation curves in Figure 3B.

Having a model that successfully captures the propagation dynamics provides evidence that the qualitative features found in the simulations are congruent with the experiments. The model shows that in the absence of positive feedback, intercellular signaling alone is not sufficient to produce super-diffusive signal propagation. To elucidate the physical mechanism responsible for the differences in signal propagation between the four strains we performed a linear analysis of the model (Supplementary). We found that in the models of the IS–PF– and IS+PF– strains signals always propagate diffusively (*∝ t*^1*/*2^). In the model IS–PF+ strain signals can propagate super-diffusively in certain specialized parameter regimes, even as the signal becomes weak due to its own diffusion. Only the IS+PF+ strain model allows for a robust, ballistic signal propagation (*∝ t*). This finding helps explain why the IS+PF+ strain exhibited much faster propagation dynamics than the other three strains in both the experiments (Fig. 3B) and numerical simulations (Fig. 3E).

Further, linear analysis of the model (Supplementary) reveals that cell growth, by increasing the amplitude of the signal profile over time, couples the propagation dynamics to the growth dynamics, leading to hyperballistic propagation in the IS+PF+ case. This finding helps explain the regime in Figure 3B and E, where the IS+PF+ strain exhibits propagation dynamics (blue) that are slightly faster than ballistic (slope greater than 1 on the log-log scale) and suggests predictions for how to enhance this regime. We investigated these and other predictions in the following section.

### Tuning the signal propagation dynamics

The PDE model makes several predictions about how the signaling dynamics can be tuned. First, the model predicts that increasing the initial concentration of erythromycin leads to a higher expression of the signal-activated regulator (either AraC or RhlI), resulting in increased sfYFP production (Fig. S5A). To test this, we performed experiments using various signal concentrations in our propagation assay. Consistent with the model’s prediction, we observed an increasing signaling distance with increasing input signal concentration, while the propagation dynamics remained ballistic regardless of the signal concentration (Fig. S5B).

Next, the linearized model predicts that signal propagation of the IS+PF+ strain can accelerate in time, provided cells are growing exponentially (Supplementary). Therefore, we investigated if extending the exponential phase of cell growth would lead to hyperballistic signal propagation. We calibrated the growth parameters of the model (*n*_0,*i*_, *g*_*i*_, *C*_*i*_, and *λ*_*i*_ in Table S2) using measured cell densities over time under different initial seeding densities (Fig. 4A). Simulations predicted that the IS+PF+ strain would show hyperballistic signal propagation at low initial cell density, whereas the IS–PF– strain, the IS+PF– strain and the IS–PF+ strain would always exhibit diffusive signal propagation (Fig. 4B,C and S6A,B).

**Figure 4:**
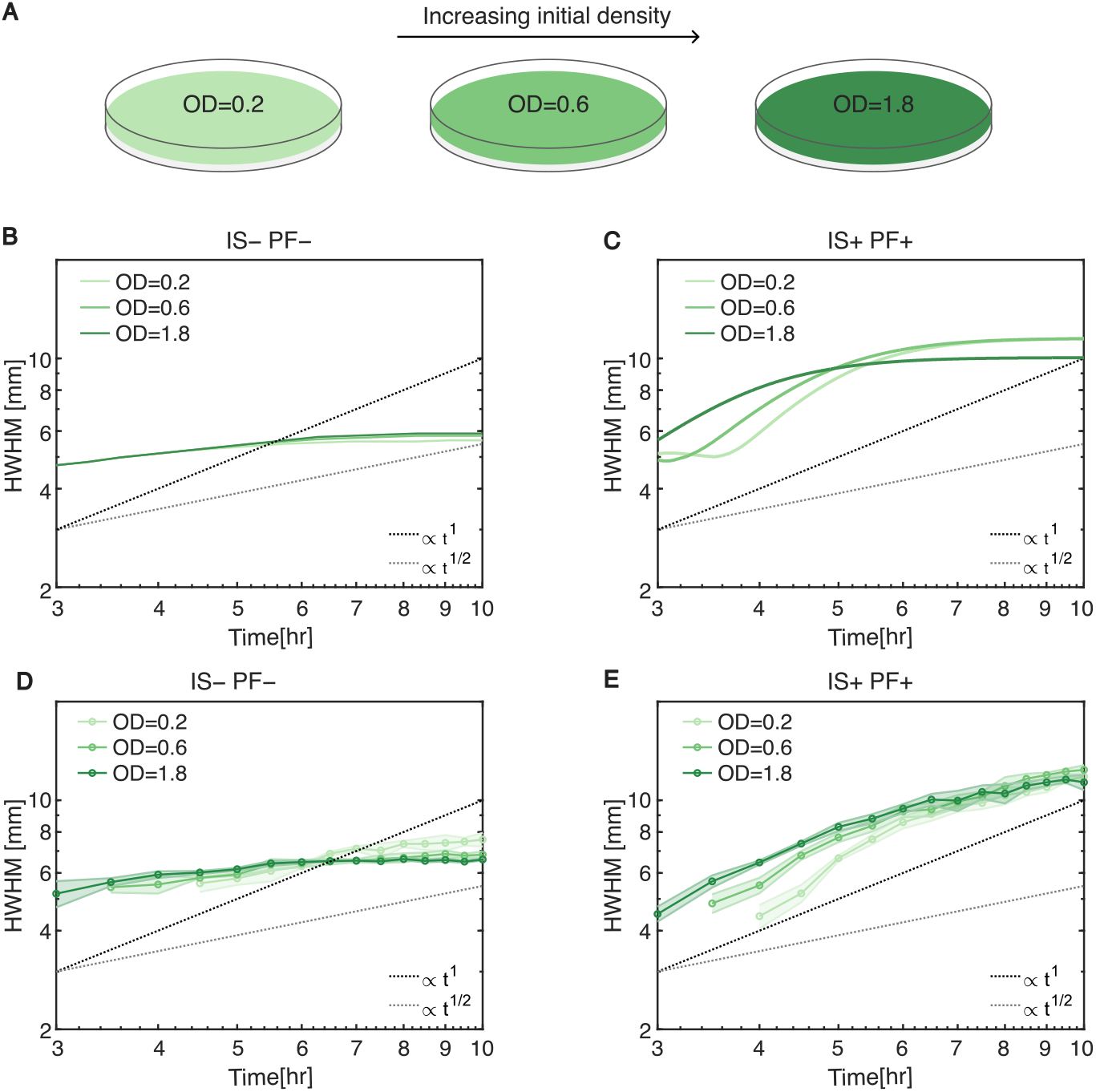
Signal propagation dynamics modulated by cell growth.(**A**) Growth modulation on an agar plate. Exponentially growing cells are plated at three initial densities (optical density = 0.2, 0.6, and 1.8), represented by three shades of green. (**B-C**)Simulated wavefront dynamics with different initial densities for the IS–PF– strain and the IS+PF+ strain. (**D-E**) Measured wavefront dynamics (HWHM) of the IS–PF– and the IS+ PF+ strain with three different initial seeding densities. Shaded region corresponds to one standard deviation of three technical replicates. Both axes are set to a logarithmic scale.

To experimentally verify these model predictions, we seeded the cells at different initial densities to modulate the growth phase of cells on agar (Fig. 4A). As the nutrient and space remained the same, populations with a smaller initial size underwent an extended exponential phase compared to populations with a larger initial size^50,51^. Growth dynamics were measured via plating assays (Methods), confirming that lower-density populations entered stationary phase roughly an hour later than higher-density populations (Fig. S3). The agar plates with lower initial density exhibited observable fluorescence signals 30 minutes to 1 hour later than plates with higher density due to microscope detection limits. Varying the initial cell density, consistent with model predictions, we found that the signaling distance and propagation rate remained similar across all initial densities in the IS–PF– strain, the IS+PF– strain and the IS-PF+ strain (Fig. 4D, S6C,D). In contrast, the IS+PF+ strain exhibited an increase of the hyperballistic signal propagation phase with decreasing OD (Fig. 4E). The enhancement is particularly pronounced with initial OD = 0.2 at 4 to 6 hours.

The IS+PF+ strain not only exhibited increased signaling distance, but also demonstrated enhanced propagation speed when coupled with exponential growth. These results confirmed that signal relay is achieved through the incorporation of both positive feedback and intercellular signaling elements, and enhanced further by cell growth.

### Robust signal propagation dynamics confirmed by absolute threshold measurements

In biological systems the half-width-half-maximum (HWHM) is not always the most meaningful measure of wavefront location. This is because the wavefront often needs to trigger a downstream component that requires a minimum concentration to activate^52^,^53^. In our system, this could correspond to the output (sfYFP) representing a transcription factor (TF) that upregulates or downregulates a downstream promoter. To account for this, we reanalyzed our data using an arbitrary threshold to measure the position of the wavefront (Fig. 5A). Specifically, we approximated fluorescence per cell by dividing total population fluorescence by population density and compared it to an arbitrary value. The wavefront position was then determined by the furthest point from the signal source at which the fluorescence per cell was greater than or equal to the threshold value. Note that the fluorescence must be scaled by the time dependent cell density in order to approximate the concentration of protein per cell.

**Figure 5:**
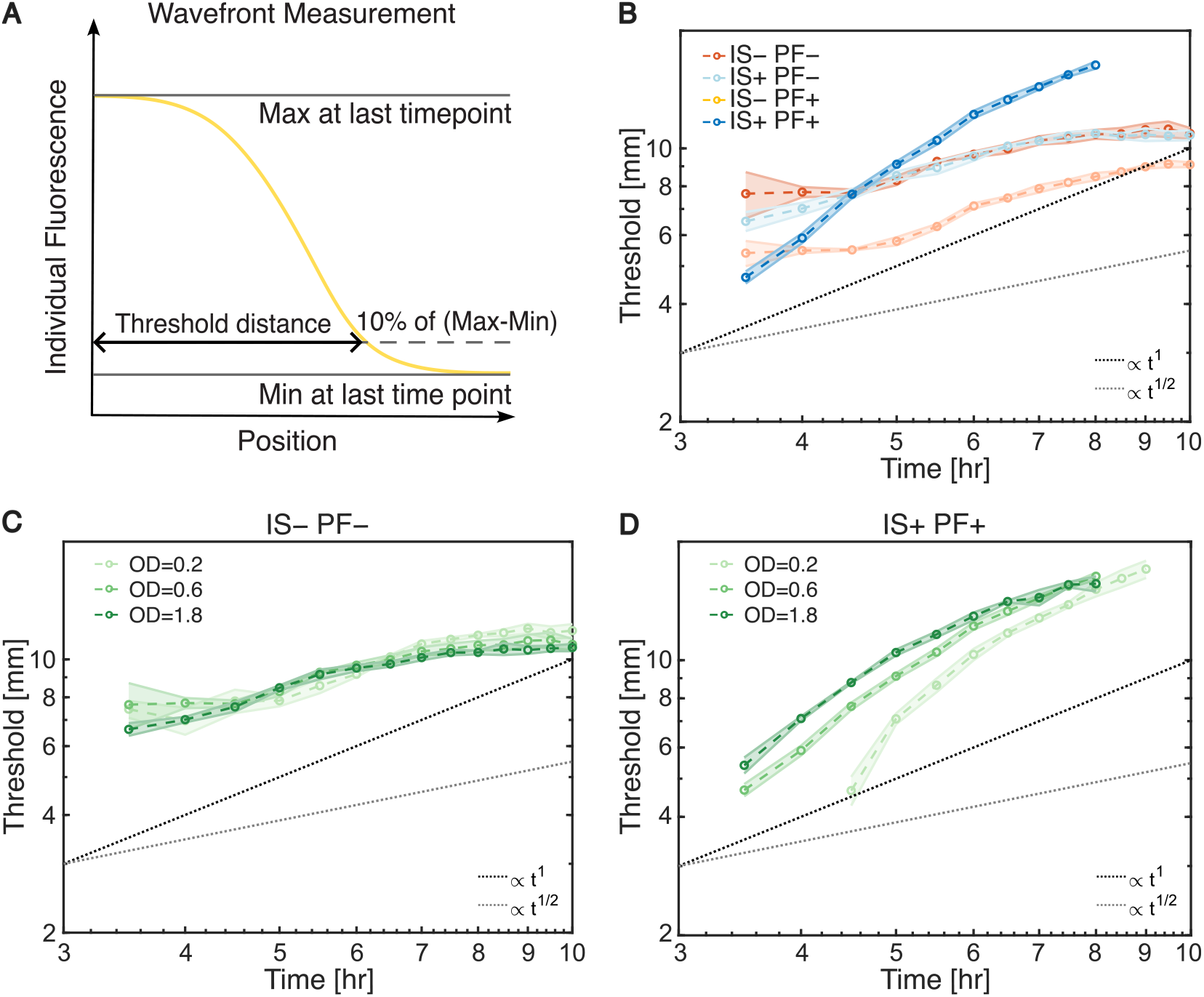
Wavefront measurement by absolute threshold confirms robust performance of growth-modulated signal relay. (**A)**Wavefront quantified using an absolute threshold of the cell-density-normalized fluorescence profile. The absolute threshold was set at 10% of the fluorescence range (max - min) at the final time point. Wavefront distance was measured based on this specific threshold. (**B)** Measured wavefront dynamics (scaled threshold) for all strains. The IS+PF+ strain exhibited super-diffusive signal propagation, while the IS–PF–, IS–PF+, IS+PF– strains showed diffusive propagation. (**C-D**) Measured wavefront dynamics(scaled threshold) of the IS–PF– and the IS+PF+ strain with three different initial seeding densities. Shaded region corresponds to one standard deviation of triplicates. Both axes are set to a logarithmic scale.

Because absolute fluorescence varies with promoter strength, RBS strength and the presence of feedback, we normalized each strain to its own maximum and used 10% of the maximum fluorescence of the corresponding strain as the threshold, so that “time-to-threshold” compares the strains on a promoter-independent scale. Similar to the wavefront measured using HWHM, the signal propagation of the IS+PF+ strain was super-diffusive, while the other three strains remained diffusive (Fig. 5B). In the context of regulating downstream promoters, the relevant TF threshold varies depending on promoter properties, such as the binding affinity of the TF to the promoter. To assess whether the observed propagation dynamics were robust to changes in threshold, we repeated the analysis using thresholds ranging from 10% to 50% of the maximum fluorescence. As expected, the wavefront distances decreased with increasing thresholds. However, the signal propagation speed of the IS+PF+ strain remained constant regardless of the threshold, while the other three strains showed decreasing speeds over time (Fig. S7).

We also asked whether increasing signal propagation observed with exponentially growing cells was also observed when defining the wavefront using specific threshold values. We reanalyzed our data with varying initial cell densities and found consistent results: the IS+PF+ strain exhibited hyperballistic signal propagation when grown at low initial density, while propagation in the IS–PF– strain, the IS–PF+ strain and the IS+PF– strain remained diffusive under all conditions (Fig. 5C-D, S8). This result further demonstrated that we can modulate the signal propagation dynamics of the IS+PF+ strain by controlling its growth dynamics.

## DISCUSSION

In this work, we engineered a synthetic version of a trigger wave network in *E. coli* that is capable of transmitting information throughout a colony at super-diffusive speeds. Like its naturally occurring counterparts in multicellular organisms, such as mitotic wave in *Xenopus laevis* egg^18^, signals in our system can travel with a constant speed. The mechanism is conceptually similar to potassium waves in *Bacillus subtilis* biofilms^22^. Unlike native bacterial QS systems, which exhibit basal auto-inducer leak and therefore act as population-density switches, our strain uses an orthogonal leak-dampener to decouple wave initiation from cell density. One reason that such small molecule trigger waves are not common in bacteria might be the metabolic burden of continuous auto-inducer synthesis and the difficulty of achieving low-leak positive feedback. By overcoming the leakiness synthetically, we show that a gene expression-based trigger wave is physically feasible.

Furthermore, we also found that in our system signals can actually accelerate and propagate hyper-ballistically. This is because the density of cells in our system can increase with time, and hence the propensity for the cells to sense and send signals at any given location can also increase. We showed that this phenomenon depends on the initial density of the cells (the longer the cells grow exponentially the faster the signal propagation), but the signalling speed might also be affected by other factors that influence growth, such as the growth medium and nutrient availability.

Of course, cellular growth has both positive and negative consequences for our system. On the plus side, growth can accelerate signaling and provide fast information transfer through the system. However, as cell density continues to increase, growth slows down and cells begin to enter stationary phase – leading to an overall slow-down of the signal as cells become less capable of responding. There are several ways in which this might be alleviated. For instance, stationary phase transcription is possible, using either phage polymerases^54,55^ or stationary phase promoters^56,57^. However, we do not currently have as many tools to regulate these systems as we do exponential phase (*σ*^70^) transcriptional systems^58,59^. Another intriguing possibility, though, is to use engineered bacteria that do not grow or divide, but are still metabolically active^60^. When used in combination with our signaling network, such “cyborg” bacteria could provide a means for long-term, and long-range responsiveness of cells to external stimuli. Alternatively, the signal relay circuit can be incorporated into bottom-up synthetic cells. These membrane-bound compartments, most commonly lipid or polymer vesicles that encapsulate a cell-free gene-expression system, have already been shown to sense, produce and exchange quorum sensing molecules^61,62^. Combining the signal relay circuit with synthetic cells would expand their utility by providing long-range communication in tissue-scale constructs such as organoids and in device-scale living materials. In parallel, this signal relay module could support patterned information processing, such as bio-computation using spatial diffusion and engineered *E*.*coli* ^63^.

It should be noted that while we embedded our engineered bacteria into agar, the same principles that guide our signalling networks should still hold for any material in which cells could be embedded that 1) supports some growth and 2) allows for extracellular signaling. Linear analysis of the signal relay circuit reveals that the wavefront speed scales with the square root of the diffusion coefficient. As a result, the super-diffusive signal propagation is largely insensitive to substrate diffusion rates, enabling a wider range of applications. Indeed, the integration of microbes with materials has given rise to a new class of engineered living materials capable of self-healing, sensing and responding to environmental signals. In these systems, diffusion-based signal propagation causes two bottlenecks: positional information is lost when signals reach distal cells only after the transcription and translation system has decayed, and the signal amplitude may drop below the activation threshold^64,65^. Incorporating our signal-relay module into cellular circuits would enable fast, efficient, and long-range signal propagation in such living materials. Dense biofilms face a similar constraint. Wild-type Bacillus subtilis in biofilms coordinates metabolism with potassium trigger waves that travel millimetres in minutes^22^. Engineered biofilms designed for self-regenerating composites or heavy-metal adsorption would likewise benefit from signal relay that is faster than diffusion, allowing distal cells to activate repair or adsorption genes while nutrients and electron donors are still available^66^. By combining positive feedback with inter-cellular signaling through quorum sensing molecules, the signal relay circuit enables centimeter scale coordination in microbial communities and can be integrated into a variety of engineered living materials or biofilms that require faster and longer-range communication.

## RESOURCE AVAILABILITY

### Lead contact

Requests for further information and resources should be directed to and will be fulfilled by the lead contact, Matthew R. Bennett.

### Materials availability

Plasmids used in this study are available through Addgene.

### Data and code availability

All data supporting the findings in the main text figures and extended data figures have been deposited in the GitHub repository at https://github.com/meidiw/Bacterial-Signal-Relay. Raw microscope images are available from the corresponding author upon reasonable request. The Matlab scripts for images analysis and model simulations can be accessed via https://github.com/meidiw/Bacterial-Signal-Relay.

## ACKNOWLEDGMENTS

This work was supported by funding from the joint National Science Foundation and National Institutes of Health Mathematical Biology Program grant 1R01GM144959 (M.W., K.J., M.R.B.), the National Science Foundation grants MCB-2118037 (L.G., S.S., A.M.), PHY-2118561 (L.G., S.S., A.M.), MCB-1936774 (M.R.B), and MCB-1936770 (K.J.), and the Welch Foundation grant C-1729 (M.R.B.). Research was sponsored by the Army Research Office and was accomplished under Cooperative Agreement Number W911NF-25-2-0076. The views and conclusions contained in this document are those of the authors and should not be interpreted as representing the official policies, either expressed or implied, of the Army Research Office or the U.S. Government. The U.S. Government is authorized to reproduce and distribute reprints for Government purposes notwithstanding any copyright notation herein.

## AUTHOR CONTRIBUTIONS

M.W., S.S., K.J., A.M. and M.R.B. conceptualized and designed the project. M.W. performed experiments and analyzed the data. L.G. and A.M. carried out the modeling and simulations. A.M., K.J. and M.R.B. oversaw the project. M.W. and L.G. drafted the manuscript, which was reviewed and revised by all authors.

## DECLARATION OF GENERATIVE AI AND AI-ASSISTED TECHNOLOGIES

The authors declare no usage of generative AI and AI-assisted technologies.

## SUPPLEMENTAL INFORMATION INDEX

Supplementary Figures S1-8 and their legends

Linear analysis of the mathematical model

Supplementary Tables S1-3 and their descriptions

## STAR METHODS

### Key Resources Table

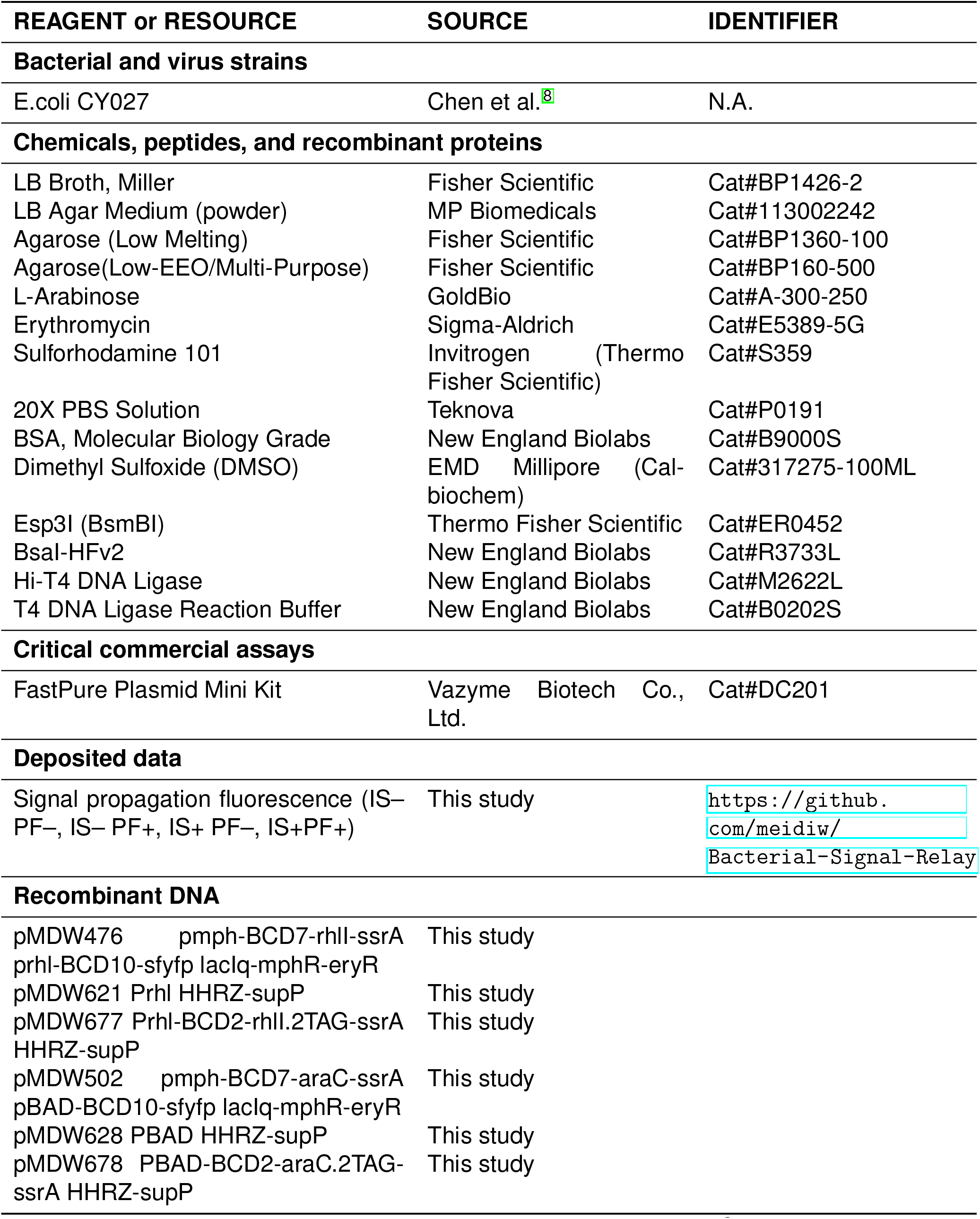
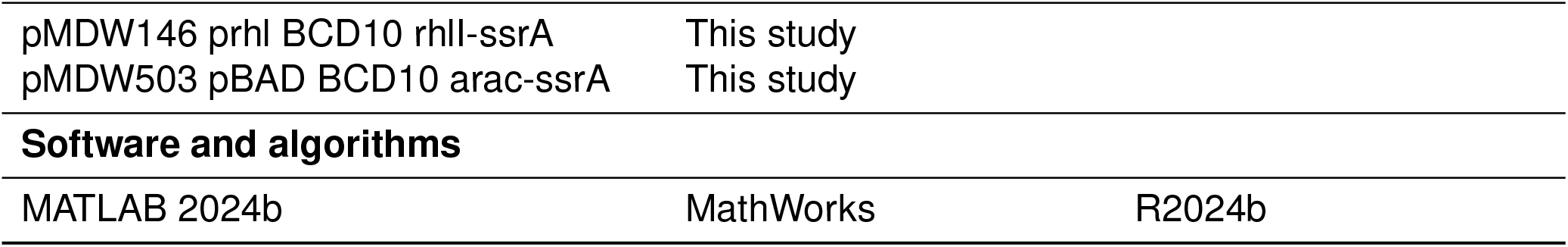

## Method details

### Plasmids and *E. coli* strain

Plasmids of all strains were constructed using golden gate cloning methods. See Table S3 for all plasmids used in this paper. All experiments were conducted in strain CY027 (*E. coli* BW25113 Δ*lacI* Δ*araC* Δ*sdiA* with constitutive *cinR* and *rhlR* integrated)^8^.

### Agar plate signal propagation assay

*E. coli* strain CY027 containing the corresponding plasmids was cultured overnight at 37°C with shaking at 250 rpm in 3 mL of LB media with kanamycin and spectinomycin (spec). The saturated culture was diluted 1:200 into 30 mL of fresh LB media containing antibiotics and incubated at 37°C with shaking at 250 rpm for less than 2 hours, until the OD_600_ reached 0.2–0.3. As cells grow, agar plates were prepared as follows: autoclaved 0.75% LB agar was melted and cooled to 55°C in a water bath. Kanamycin, spectinomycin, and arabinose were added to the agar to final concentrations of 50 mg/L, 50 mg/L, and 0.1% (w/w), respectively. Then, 3.5 mL of the well-mixed agar was aliquoted into each 60 mm petri dish to form an even layer. Separately, 0.55% low-melt LB agar was melted, supplemented with kanamycin, spectinomycin, and arabinose, aliquoted into small tubes and kept warm in a 42°C water bath. Early exponential-phase cells were concentrated and mixed with the 0.55% low-melt LB agar to achieve a final OD_600_ of 0.6 in all experiments, unless otherwise specified. Next, 500 *μ*L of the cell-agar mixture was aliquoted onto the prepared petri dish containing the 0.75% LB agar layer. Each condition included three technical replicates. All plates were allowed to solidify at room temperature for 10 minutes. Next, 2 *μ*L of a solution containing erythromycin (2 mg/mL) and the red fluorescent dye sulforhodamine 101 (10 *μ*g/mL) was deposited at the center of each plate. The plates were incubated at 37°C for 10 hours. Imaging was conducted every 30 minutes under brightfield, yellow fluorescence, and red fluorescence using a Nikon fluorescence stereomicroscope from 3 to 10 hours.

### Fluorescence image analysis

We compiled the yellow fluorescence images at different time points to create the images shown in Fig. 3A. Data analysis to measure the wavefront movement of each strain was performed with custom MATLAB code. This code determined the regions of interest (ROIs) using the red fluorescence images, which is a rectangle mask that is around 1000 pixels long from the center of the signal source to the edge of the field of view and 40 pixels wide. The radial yellow fluorescence profile at each time point was calculated by averaging the fluorescence along the width and subtracting the basal fluorescence intensity from the autofluorescence of E.coli and the LB media. This profile was noise-filtered by applying the Savitzky-Golay filter. The wavefront movement was calculated in two ways. The first method uses the half-width of a line shape at half of its maximum amplitude (HWHM) to represent the wavefront. The second method calculated the wavefront by normalizing the fluorescence profile by cell density and measuring the width of a line shape at a specific threshold *L*_*th*_. The cell density was estimated from the growth dynamics experiment on agar plates. *L*_*th*_, set as 0.1 *\* L*_*max*_ in Fig. 5, is determined by the maximum amplitude (*L*_*max*_) of the circuit at the end of the experiment.

All fluorescence images are flat-field corrected to compensate for uneven illumination and detection. Uniform fluorescence samples were prepared by mixing fluorescein with melted 1.5% LB agar to final concentrations of 5 nM, 10 nM and 15 nM to perform flat-field correction of different fluorescence. *S*_*ij*_ represents fluorescence intensity of the original image of the uniform sample at position (i.j). *S*^*′*^ represents the average intensity of the original uniform sample. For each *I*_*ij*_, original fluorescence intensity of the experiment sample, the flat-field corrected 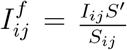.

### Self-activation measurement on agar

*E. coli* strain CY027 containing the corresponding plasmids were cultured overnight. Plates were prepared using the same method as described in the signal propagation assay. Each strain had three technical replicates. All plates were allowed to solidify at room temperature for 10 minutes and incubated at 37°C. We imaged each plate under brightfield and yellow fluorescence at 10 hours.

### 96-well plate assays

To measure the response of each strain to erythromycin, a single colony of each strain was inoculated into 3 mL of LB medium containing kanamycin and spectinomycin and grown for 12–16 hours. The saturated cultures were diluted 1:100 into fresh LB medium with antibiotics and aliquoted into a 96-well plate. The TECAN Spark plate reader was preheated to 37^°^C, and the plate was placed into a small humidity cassette containing 4 mL of water on each side to minimize evaporation. The following settings were used for the TECAN Spark plate reader: temperature set to 37^°^C, kinetic loop for 11 hours, orbital shaking at 510 rpm, and event acquisition every 10 minutes. After shaking for 1 hour, the program was paused, and 100 *μ*L of media containing 2X inducer was added to the cultures. The program then resumed. The final erythromycin concentrations tested were 100 *μ*g/mL, 30 *μ*g/mL, 10 *μ*g/mL, 3 *μ*g/mL, 1 *μ*g/mL, 0.3 *μ*g/mL, 0.1 *μ*g/mL, and 0 *μ*g/mL. The response curves shown in Figure S1a were obtained after 11 hours, when the cells reached the stationary phase.

### Growth dynamics on agar plates

To measure the growth dynamics of each circuit on agar plates, we developed a sample and plating assay to count the colonies. Plates were prepared using the same method as described in the signal propagation assay. We use 1000 *μ*L pipette tips to collect three samples from the plate every hour for a total of 6 hours. The samples were immediately mixed with PBS buffer to halt growth and vigorously vortexed for 5 minutes before being serially diluted to the appropriate cell density. The dilution concentration was determined through tests to keep the cell densities between 20 and 150 per sample, ensuring that the colonies spread sufficiently for accurate counting. For each sample, 10 *μ*L of diluted cell culture was spread on a LB agar plate with three replicates. The averaged cell count of the three replicates represents the cell count of the sample. This process was repeated for each circuit at three different initial densities, with three technical replicates.

The growth parameters of each circuit were estimated using MATLAB. We modeled the cell density *n*_*i*_(*t*) with the following equation proposed by Huang^49^:

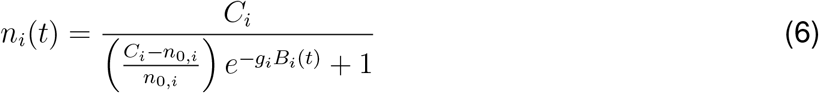

where *n*_0,*i*_ is the initial cell number and

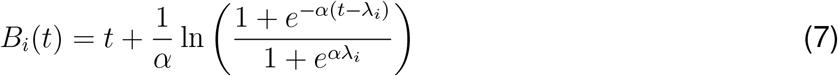

quantifies the cell growth from an initial lag phase with lag time *λ*_*i*_. The average cell density we acquired from the experiment is used to estimate the growth parameters for each circuit with each initial cell density. The experimental data and fitted growth curve is shown in Figure S3. The estimate growth parameters from these experiments are shown in Table S2.

### Cellular growth function

In this section, we define the cell growth function *f*_*i*_(*t*) used in equation (2) for each circuit with index *i*. Multiple studies have previously established that the transcriptional rate increases sub-linearly with growth rate^33,67–69^. We therefore estimate the transcriptional rate as proportional to 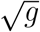 as it follows their measured data closely without introducing extra parameters. This growth envelope function restricts protein production to the period of significant cell growth, after the lag phase and before stationary phase (Fig. S4). From equation(4), we define a new dimensionless parameter *ρ*_*i*_ *≡ n*_0,*i*_*/C*_*i*_. Normalizing *n*_*i*_(*t*) by the circuit-specific carrying capacity *C*_*i*_, we arrive at

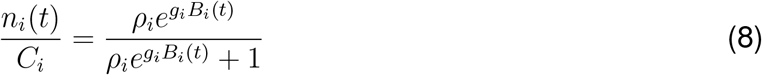

for *ρ*_*i*_ *<<* 1. Now we take the time derivative of the previous equation to get

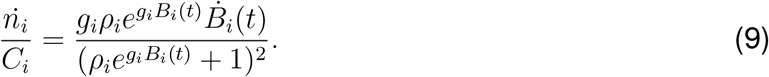

In our model, the growth rate is 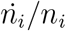, therefore the transcription rate per cell is 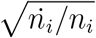. The total transcription rate for *n*_*i*_ cells at a particular location is thus proportional to 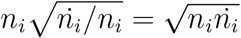

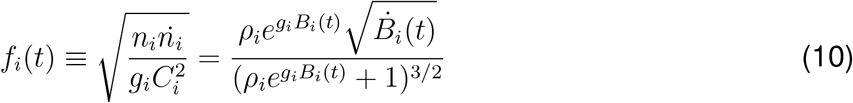

where 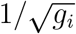 is introduced in the final step to make the resulting expression dimensionless.

### Determination of parameters from liquid culture data

The hill coefficient (*H*_1_ and *H*_3_) and half-maximal concentration (*S*_1_ and *S*_3_) describe intracellular binding events and are therefore expected to remain unchanged between liquid and solid LB. To determine particular parameters for the model, we use fitted data taken from the experiments done in liquid culture. To do so, we first neglect the spatial effects of diffusion for a well mixed medium. Second, we also note that the four circuits can be categorized in two distinct groups: the IS-PF+ case is simply the addition of a responder to the control circuit and likewise for the IS+PF+ case being the quorum sensing circuit with the responder added. Hence, we can assume that in a well mixed medium, both the quorum sensing and control circuits are identical. Thus we simply write

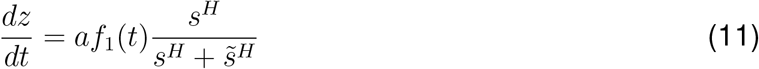

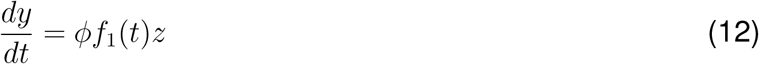

for the control and quorum sensing circuits where, for the sake of our argument, we have temporarily dropped the subscript labeling for each circuit and likewise temporarily renamed *s*_*i*_ as 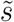. The source term *s* is some constant due to the medium. We solve the ordinary differential equation for *z*(*r, t*):

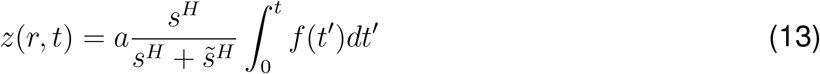

where we use a placeholder variable *t*^′^ for time *t*. The solution of *y*(*r, t*) can now be rewritten as

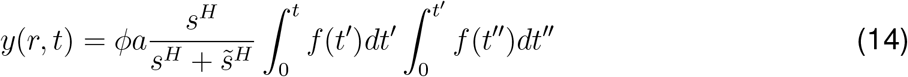

where we once again use a placeholder variable *t*^″^ for time. At long times *y*_*∞*_, the integrals become definite and collapse into a constant

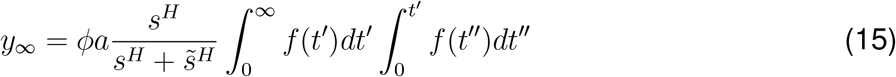

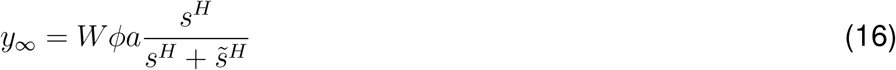

where *W* is some constant. Equation (16) indicates YFP at long times depends on *s* only via the activation function, implying that the liquid culture parameters *s*_1_ and *H*_1_ can be directly taken from the fitted data. A similar argument is made for the quorum sensing circuit as *z* is uniform in liquid culture. Hence, from Figure S1a: *H*_1_ = 1.6, *s*_1_ = 8.7 *μ*g*/*mL, *H*_3_ = 1.9, and *s*_3_ = 6.3 *μ*g*/*mL. This leaves *J*_2_, *b*_2_, *z*_2_, *J*_4_, *b*_4_, and *z*_4_ as free parameters. Our choice of parameters is further explained in the following section.

### Choice of other parameters

We approximate the droplet size to be *R*_0_ = 0.5 mm. The diffusion constant of an erythromycin molecule was estimated to be *D* = 500 *μ*m^2^*/*s. For the secondary diffusing molecule C4-HSL *z*, we used the fact that the molecular weight of erythromycin (733 g*/*mol) is about 1.6 times larger than that of C4-HSL (171 g*/*mol) to set the diffusivity *E ≈* 1.6 *· D* = 800 *μ*m^2^*/*s. We set the initial concentration as *s*_0_ = 2000 *μ*g*/*mL as in the experiments.

Starting with the feedback only case (IS-PF+), *b*_2_ qualitatively amplifies the strength of the of response in rhlI once the concentration threshold, *z*_2_, is met. Hence, an increase in *b*_2_ changes the propagation of the feedback circuit from to super-diffusive, an outcome not seen in the experiments. (Fig. S9A) By changing *z*_2_, we likewise change the time in which the threshold is met (Fig. S9B). Lastly, *J*_*i*_ is the hill coefficient for rhlI production. In this case, *J*_2_ sets the sharpness of the response once the threshold has been met (Fig. S9C). A higher *J*_2_ corresponds to a faster propagation, if the feedback begins sooner (*b*_2_ gets smaller) and/or the threshold concentration is smaller (*z*_2_ gets smaller). We set *b*_2_ = 3.5, *J*_2_ = 2, and *z*_2_ = 0.5 *μ*g*/*mL to approximate the experimental results in Fig. 3.

For the IS+PF+ circuit, *b*_4_, *z*_4_, and *J*_4_ behave qualitatively the same as previously discussed. Calibration of each parameter value was taken by approximating the experimental amplitude of the IS+PF+ circuit. Thus, we set *b*_4_ = 3.5, *z*_4_ = 0.05 *μ*g*/*mL and *J*_4_ = 2 (Fig. S9D-F).

### Effects of AraC degradation in signal propagation

To explore the effects of the half-maximal concentration *z*_2_ and *z*_4_ for both the IS-PF+ and IS+PF+ cases, we incorporate the effects of degradation of AraC in the positive feedback circuit. The new model with AraC degradation is

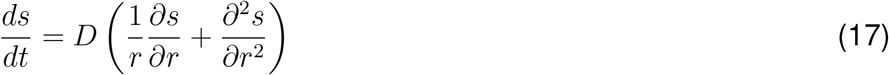

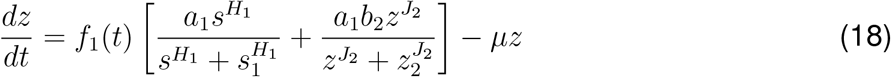

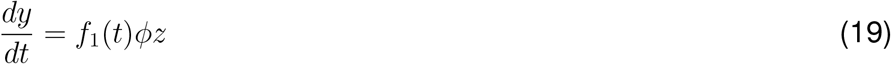

for the IS-PF+ strain. Here, *μ* = 10 h^−1^ is the degradation rate. Figure S10 shows the results of having *z*_2_ = *z*_4_ with AraC degradation. When degradation is included, the output of the model is similar to that of Figure 3 in the main text. Thus, we can assume that our choice of *z*_2_ *> z*_4_ is consistent with the fact that the responder protein AraC also degrades as it accumulates in the circuit.

## References

1. Li, P., and Elowitz, M.B. (2019). Communication codes in developmental signaling pathways. Development 146, dev170977.

2. Zinner, M., Lukonin, I., and Liberali, P. (2020). Design principles of tissue organisation: How single cells coordinate across scales. Current Opinion in Cell Biology 67, 37–45.

3. Whiteley, M., Diggle, S.P., and Greenberg, E.P. (2017). Progress in and promise of bacterial quorum sensing research. Nature 551, 313–320.

4. Bassler, B.L. (2002). Small talk: cell-to-cell communication in bacteria. Cell 109, 421–424.

5. Yashiroda, Y., and Yoshida, M. (2019). Intraspecies cell–cell communication in yeast. FEMS Yeast Research 19, foz071.

6. Wu, F., Ha, Y., Weiss, A., Wang, M., Letourneau, J., Wang, S., Luo, N., Huang, S., Lee, C.T., David, L.A. et al. (2022). Modulation of microbial community dynamics by spatial partitioning. Nature Chemical Biology 18, 394–402.

7. Basu, S., Gerchman, Y., Collins, C.H., Arnold, F.H., and Weiss, R. (2005). A synthetic multicellular system for programmed pattern formation. Nature 434, 1130–1134.

8. Chen, Y., Kim, J.K., Hirning, A.J., Josić, K., and Bennett, M.R. (2015). Emergent genetic oscillations in a synthetic microbial consortium. Science 349, 986–989.

9. Kong, W., Meldgin, D.R., Collins, J.J., and Lu, T. (2018). Designing microbial consortia with defined social interactions. Nature Chemical Biology 14, 821–829.

10. Alnahhas, R.N., Sadeghpour, M., Chen, Y., Frey, A.A., Ott, W., Josić, K., and Bennett, M.R. (2020). Majority sensing in synthetic microbial consortia. Nature Communications 11, 3659.

11. Combarnous, Y., and Nguyen, T.M.D. (2020). Cell communications among microorganisms, plants, and animals: origin, evolution, and interplays. International Journal of Molecular Sciences 21, 8052.

12. Mukherjee, S., and Bassler, B.L. (2019). Bacterial quorum sensing in complex and dynamically changing environments. Nature Reviews Microbiology 17, 371–382.

13. Pesci, E.C., Milbank, J.B., Pearson, J.P., McKnight, S., Kende, A.S., Greenberg, E.P., and Iglewski, B.H. (1999). Quinolone signaling in the cell-to-cell communication system of pseudomonas aeruginosa. Proceedings of the National Academy of Sciences 96, 11229–11234.

14. Arora, D.P., Hossain, S., Xu, Y., and Boon, E.M. (2015). Nitric oxide regulation of bacterial biofilms. Biochemistry 54, 3717–3728.

15. Cao, Y., Ryser, M.D., Payne, S., Li, B., Rao, C.V., and You, L. (2016). Collective space-sensing coordinates pattern scaling in engineered bacteria. Cell 165, 620–630.

16. Kim, J.K., Chen, Y., Hirning, A.J., Alnahhas, R.N., Josić, K., and Bennett, M.R. (2019). Long-range temporal coordination of gene expression in synthetic microbial consortia. Nature chemical biology 15, 1102–1109.

17. Gelens, L., Anderson, G.A., and Ferrell Jr, J.E. (2014). Spatial trigger waves: positive feed-back gets you a long way. MBoC 25, 3486–3493.

18. Chang, J.B., and Ferrell Jr, J.E. (2013). Mitotic trigger waves and the spatial coordination of the xenopus cell cycle. Nature 500, 603–607.

19. Co, H.K., Wu, C.C., Lee, Y.C., and Chen, S.h. (2024). Emergence of large-scale cell death through ferroptotic trigger waves. Nature pp. 1–9.

20. Dieterle, P.B., Min, J., Irimia, D., and Amir, A. (2020). Dynamics of diffusive cell signaling relays. Elife 9, e61771.

21. Patel, K., Rodriguez, C., Stabb, E.V., and Hagen, S.J. (2021). Wavelike propagation of quorum activation through a spatially distributed bacterial population under natural regulation. Phys. Biol. 18, 046008.

22. Prindle, A., Liu, J., Asally, M., Ly, S., Garcia-Ojalvo, J., and Süel, G.M. (2015). Ion channels enable electrical communication in bacterial communities. Nature 527, 59–63.

23. Ward, J.P., King, J.R., Koerber, A., Williams, P., Croft, J., and Sockett, R. (2001). Mathematical modelling of quorum sensing in bacteria. Mathematical Medicine and Biology 18, 263–292.

24. Kim, M.K., Ingremeau, F., Zhao, A., Bassler, B.L., and Stone, H.A. (2016). Local and global consequences of flow on bacterial quorum sensing. Nature Microbiology 1, 1–5.

25. Darch, S.E., Simoska, O., Fitzpatrick, M., Barraza, J.P., Stevenson, K.J., Bonnecaze, R.T., Shear, J.B., and Whiteley, M. (2018). Spatial determinants of quorum signaling in a pseudomonas aeruginosa infection model. Proceedings of the National Academy of Sciences 115, 4779–4784.

26. Papenfort, K., and Bassler, B.L. (2016). Quorum sensing signal–response systems in gram-negative bacteria. Nature Reviews Microbiology 14, 576–588.

27. Dalwadi, M.P., and Pearce, P. (2021). Emergent robustness of bacterial quorum sensing in fluid flow. Proceedings of the National Academy of Sciences 118, e2022312118.

28. Showalter, K., and Tyson, J.J. (1987). Luther’s 1906 discovery and analysis of chemical waves. Journal of Chemical Education 64, 742.

29. Tyson, J.J., and Keener, J.P. (1988). Singular perturbation theory of traveling waves in excitable media (a review). Physica D: Nonlinear Phenomena 32, 327–361.

30. Dilanji, G.E., Langebrake, J.B., De Leenheer, P., and Hagen, S.J. (2012). Quorum activation at a distance: spatiotemporal patterns of gene regulation from diffusion of an autoinducer signal. Journal of the American Chemical Society 134, 5618–5626.

31. Patel, K., Rodriguez, C., Stabb, E.V., and Hagen, S.J. (2020). Spatially propagating activation of quorum sensing in vibrio fischeri and the transition to low population density. Phys. Rev. E 101, 062421.

32. Danino, T., Mondragón-Palomino, O., Tsimring, L., and Hasty, J. (2010). A synchronized quorum of genetic clocks. Nature 463, 326–330.

33. Klumpp, S., Zhang, Z., and Hwa, T. (2009). Growth rate-dependent global effects on gene expression in bacteria. Cell 139, 1366–1375.

34. Scott, M., Gunderson, C.W., Mateescu, E.M., Zhang, Z., and Hwa, T. (2010). Interdependence of cell growth and gene expression: origins and consequences. Science 330, 1099– 1102.

35. Meyer, A.J., Segall-Shapiro, T.H., Glassey, E., Zhang, J., and Voigt, C.A. (2019). Escherichia coli “marionette” strains with 12 highly optimized small-molecule sensors. Nature Chemical Biology 15, 196–204.

36. Pesci, E.C., Pearson, J.P., Seed, P.C., and Iglewski, B.H. (1997). Regulation of las and rhl quorum sensing in pseudomonas aeruginosa. Journal of Bacteriology 179, 3127–3132.

37. Ho, J.M., Miller, C.A., Parks, S.E., Mattia, J.R., and Bennett, M.R. (2021). A suppressor trnamediated feedforward loop eliminates leaky gene expression in bacteria. Nucleic Acids Res. 49, e25–e25.

38. Andersen, J.B., Sternberg, C., Poulsen, L.K., Bjørn, S.P., Givskov, M., and Molin, S. (1998). New unstable variants of green fluorescent protein for studies of transient gene expression in bacteria. Applied and Environmental Microbiology 64, 2240–2246.

39. Pletnev, P., Osterman, I., Sergiev, P., Bogdanov, A., and Dontsova, O. (2015). Survival guide: Escherichia coli in the stationary phase. Acta Naturae () 7, 22–33.

40. Navarro Llorens, J.M., Tormo, A., and Martínez-García, E. (2010). Stationary phase in gram-negative bacteria. FEMS Microbiology Reviews 34, 476–495.

41. Schleif, R. (2000). Regulation of the l-arabinose operon of escherichia coli. Trends in Genetics 16, 559–565.

42. Gao, M., Zheng, H., Ren, Y., Lou, R., Wu, F., Yu, W., Liu, X., and Ma, X. (2016). A crucial role for spatial distribution in bacterial quorum sensing. Scientific Reports 6, 34695.

43. Alberghini, S., Polone, E., Corich, V., Carlot, M., Seno, F., Trovato, A., and Squartini, A. (2009). Consequences of relative cellular positioning on quorum sensing and bacterial cellto-cell communication. FEMS Microbiology Letters 292, 149–161.

44. Pearson, J.P., Van Delden, C., and Iglewski, B.H. (1999). Active efflux and diffusion are involved in transport of pseudomonas aeruginosa cell-to-cell signals. Journal of bacteriology 181, 1203–1210.

45. Pai, A., and You, L. (2009). Optimal tuning of bacterial sensing potential. Molecular Systems Biology 5, 286.

46. Glass, D.S., Jin, X., and Riedel-Kruse, I.H. (2021). Nonlinear delay differential equations and their application to modeling biological network motifs. Nature Communications 12, 1788.

47. Rombouts, J., Gelens, L., and Erneux, T. (2019). Travelling fronts in time-delayed reaction– diffusion systems. Philosophical Transactions of the Royal Society A 377, 20180127.

48. Moreira Gomes, J., Lobosco, M., Weber dos Santos, R., and Cherry, E.M. (2019). Delay differential equation-based models of cardiac tissue: Efficient implementation and effects on spiral-wave dynamics. Chaos: An Interdisciplinary Journal of Nonlinear Science 29.

49. Huang, L. (2011). A new mechanistic growth model for simultaneous determination of lag phase duration and exponential growth rate and a new be? lehdrádek-type model for evaluating the effect of temperature on growth rate. Food Microbiology 28, 770–776.

50. Cooper, A., Dean, A., and Hinshelwood, C.N. (1968). Factors affecting the growth of bacterial colonies on agar plates. Proceedings of the Royal Society of London. Series B. Biological Sciences 171, 175–199.

51. Shao, X., Mugler, A., Kim, J., Jeong, H.J., Levin, B.R., and Nemenman, I. (2017). Growth of bacteria in 3-d colonies. PLoS Computational Biology 13, e1005679.

52. Landry, B.P., Palanki, R., Dyulgyarov, N., Hartsough, L.A., and Tabor, J.J. (2018). Phosphatase activity tunes two-component system sensor detection threshold. Nature Communications 9, 1433.

53. Todeschini, A.L., Georges, A., and Veitia, R.A. (2014). Transcription factors: specific dna binding and specific gene regulation. Trends in Genetics 30, 211–219.

54. Studier, F.W., and Moffatt, B.A. (1986). Use of bacteriophage t7 rna polymerase to direct selective high-level expression of cloned genes. Journal of Molecular Biology 189, 113–130.

55. Tabib-Salazar, A., Liu, B., Barker, D., Burchell, L., Qimron, U., Matthews, S.J., and Wigneshweraraj, S. (2018). T7 phage factor required for managing rpos in escherichia coli. Proceedings of the National Academy of Sciences 115, E5353–E5362.

56. Hengge-Aronis, R. (2002). Signal transduction and regulatory mechanisms involved in control of the s (rpos) subunit of rna polymerase. Microbiology and Molecular Biology Reviews 66, 373–395.

57. Lacour, S., and Landini, P. (2004). s-dependent gene expression at the onset of stationary phase in escherichia coli: function of s-dependent genes and identification of their promoter sequences. Journal of Bacteriology 186, 7186–7195.

58. Shimada, T., Makinoshima, H., Ogawa, Y., Miki, T., Maeda, M., and Ishihama, A. (2004). Classification and strength measurement of stationary-phase promoters by use of a newly developed promoter cloning vector. Journal of Bacteriology 186, 7112–7122.

59. Tripathi, L., Zhang, Y., and Lin, Z. (2014). Bacterial sigma factors as targets for engineered or synthetic transcriptional control. Frontiers in Bioengineering and Biotechnology 2, 33.

60. Baghdasaryan, O., Lee-Kin, J., and Tan, C. (2024). Architectural engineering of cyborg bacteria with intracellular hydrogel. Materials Today Bio 28, 101226.

61. Lentini, R., Martín, N.Y., Forlin, M., Belmonte, L., Fontana, J., Cornella, M., Martini, L., Tamburini, S., Bentley, W.E., Jousson, O. et al. (2017). Two-way chemical communication between artificial and natural cells. ACS Central Science 3, 117–123.

62. Smith, J.M., Hartmann, D., and Booth, M.J. (2023). Engineering cellular communication between light-activated synthetic cells and bacteria. Nature Chemical Biology 19, 1138– 1146.

63. Fedorec, A.J., Treloar, N.J., Wen, K.Y., Dekker, L., Ong, Q.H., Jurkeviciute, G., Lyu, E., Rutter, J.W., Zhang, K.J., Rosa, L. et al. (2024). Emergent digital bio-computation through spatial diffusion and engineered bacteria. Nature Communications 15, 4896.

64. Molinari, S., Tesoriero, R.F., and Ajo-Franklin, C.M. (2021). Bottom-up approaches to engineered living materials: Challenges and future directions. Matter 4, 3095–3120.

65. Dupin, A., Aufinger, L., Styazhkin, I., Rothfischer, F., Kaufmann, B.K., Schwarz, S., Galensowske, N., Clausen-Schaumann, H., and Simmel, F.C. (2022). Synthetic cell–based materials extract positional information from morphogen gradients. Science Advances 8, eabl9228.

66. Zhu, X., Xiang, Q., Chen, L., Chen, J., Wang, L., Jiang, N., Hao, X., Zhang, H., Wang, X., Li, Y. et al. (2024). Engineered bacillus subtilis biofilm@ biochar living materials for in-situ sensing and bioremediation of heavy metal ions pollution. Journal of Hazardous Materials 465, 133119.

67. Liang, S.T., Bipatnath, M., Xu, Y.C., Chen, S.L., Dennis, P., Ehrenberg, M., and Bremer, H. (1999). Activities of constitutive promoters in escherichia coli. Journal of Molecular Biology 292, 19–37.

68. Bremer, H., and Dennis, P.P. (2008). Modulation of chemical composition and other parameters of the cell at different exponential growth rates. Ecosal Plus 3, 10–1128.

69. Calabrese, L., Ciandrini, L., and Cosentino Lagomarsino, M. (2024). How total mrna influences cell growth. Proceedings of the National Academy of Sciences 121, e2400679121.

