## Supplementary for "Fast, long-range intercellular signal propagation through growth assisted positive feedback"

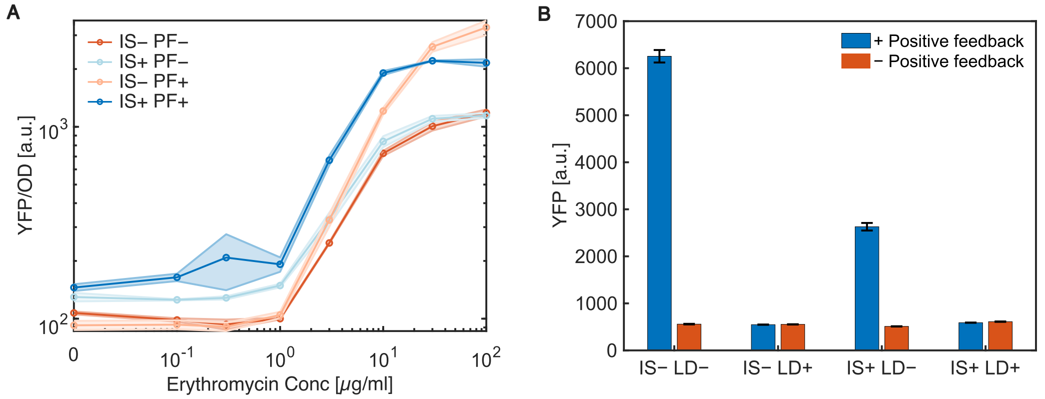


Figure S1: Characterization of the engineered strains. (**A**) Fluorescence intensity of IS–PF–, IS–PF+, IS+PF–, and IS+PF+ strains in liquid culture with varying erythromycin concentrations. The IS–PF+ strain and the IS+PF+ strain show increased fluorescence with higher erythromycin concentrations, confirming effective positive feedback without self-activation. (**B**) Self-activation was measured in the IS–PF+ and the IS+PF+ strains in the absence of erythromycin, both with and without the leak dampener module. Fluorescence levels indicate that the dampener effectively prevents self-activation.

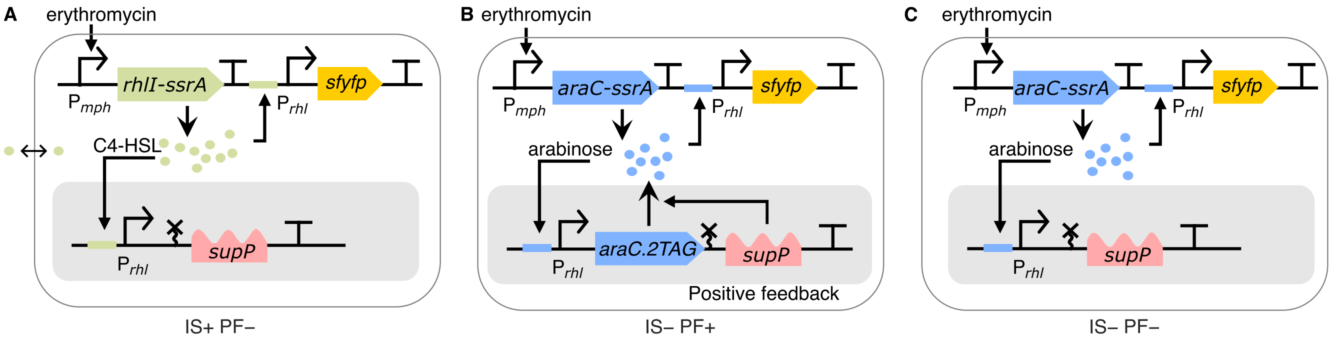


Figure S2: Circuit diagram of the strains without intercellular signaling or positive feedback or both. (**A**) Circuit diagram of the IS+PF– strain. (**B**)Circuit diagram of the IS–PF+ strain. (**C**)Circuit diagram of the IS–PF– strain.

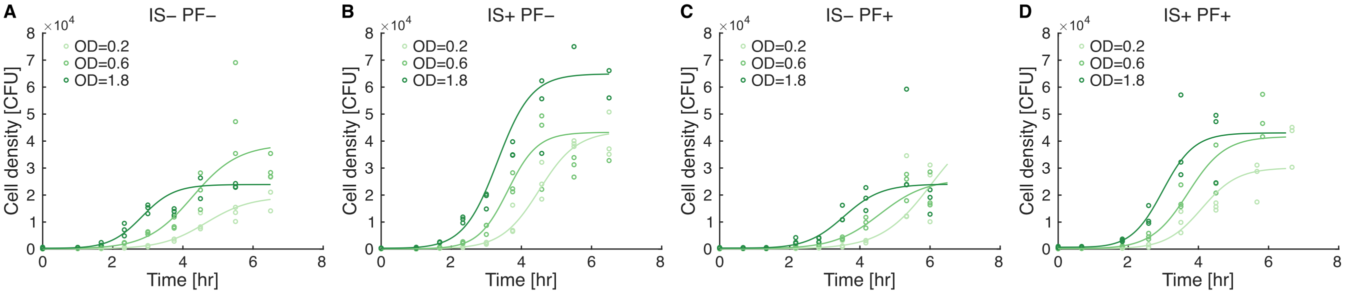


Figure S3: Cell density over time for strains embedded in soft agar. (**A**) IS–PF– strain, (**B**) IS+PF– strain, (**C**) IS–PF+ strain, and (**D**) IS+PF– strain. Cells are seeded at three initial ODs: 0.2, 0.6 and 1.8. Three samples were taken at each time point for each strain. Data show the growth transition from lag phase to exponential phase to stationary phase.

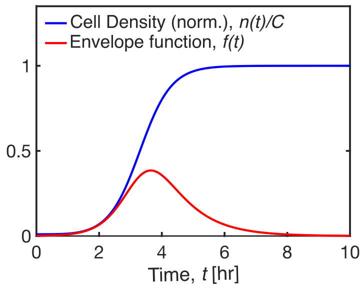


Figure S4: Example of envelope function f(t) corresponding to cell density curve n(t) with parameters $g = 2/h$, $n_0/C = 0.01$, $\lambda = 1h$, and $\alpha = 4/h$.

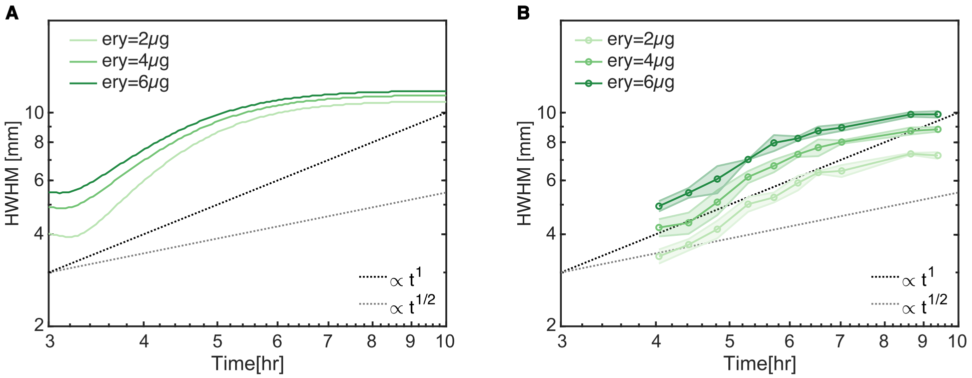


Figure S5: Effect of initial erythromycin concentration on signaling distance. (**A**) Simulation results predict an increase in signaling distance with higher initial concentrations of erythromycin (**B**) Experiment results with varying concentrations of erythromycin, consistent with model predictions.

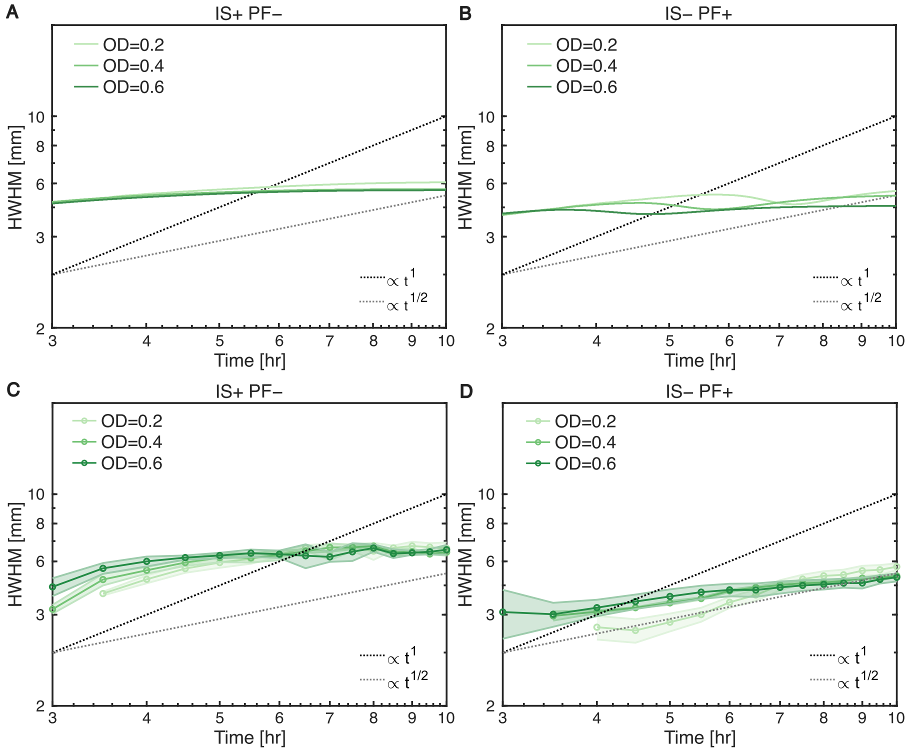


Figure S6: Growth modulated wavefront propagation (HWHM) of the IS+PF– and the IS–PF+ strain (**A-B**)Simulated wavefront dynamics with three different initial seeding densities. (**C-D**) Measured wavefront dynsmics with three different initial seeding densities (optical density = 0.2, 0.6 and 1.8). Shaded region corresponds to one standard deviation of three technical replicates. Both axes are set to a logarithmic scale.

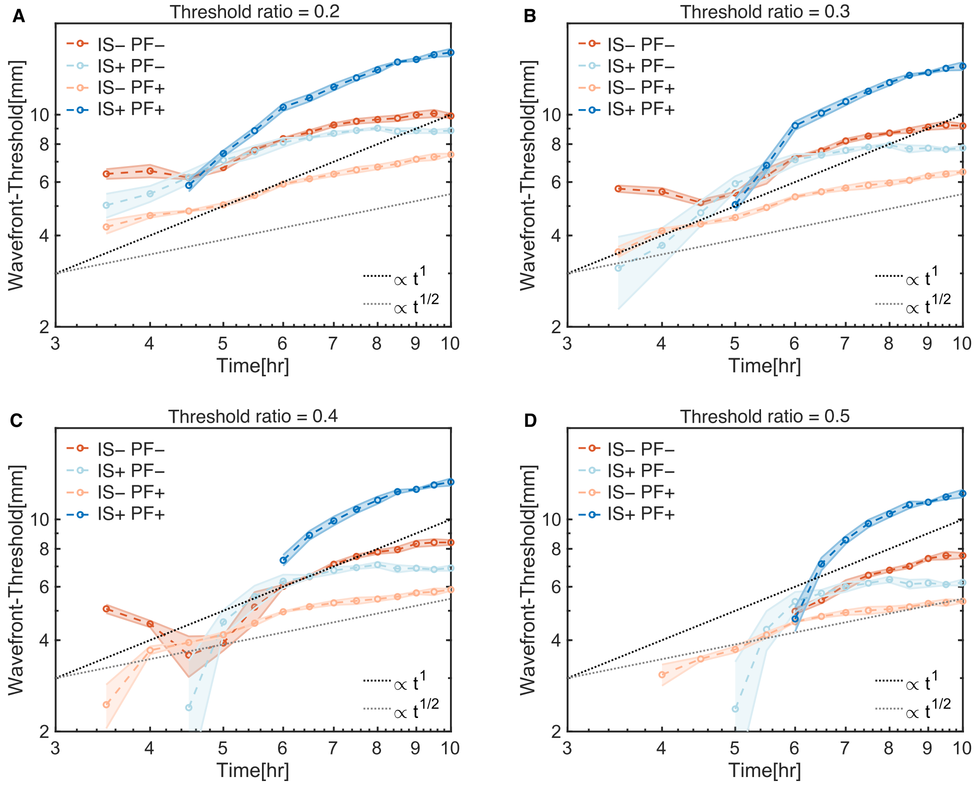


Figure S7: Measured wavefront propagation (scaled threshold) using different threshold. (**A**) $L_{th} = 0.2 * L_{max}$. (**B**) $L_{th} = 0.3 * L_{max}$. (**C**) $L_{th} = 0.4 * L_{max}$. (**D**) $L_{th} = 0.5 * L_{max}$.

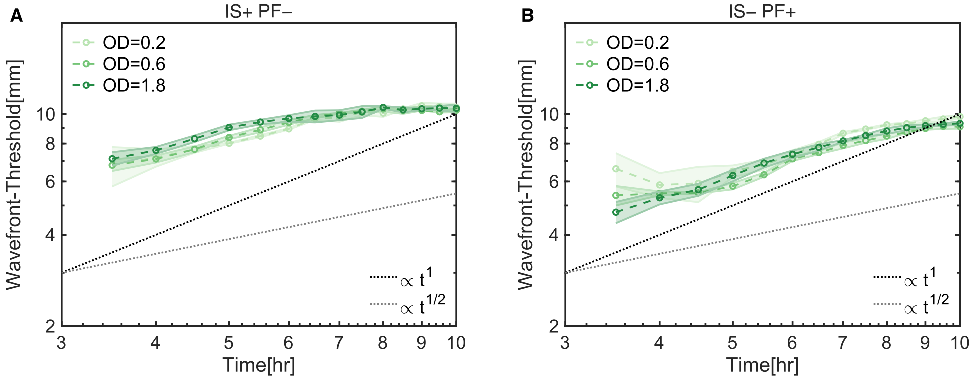


Figure S8: Growth modulated wavefront propagation (scaled threshold). (**A**) The IS+PF– strain with three different initial seeding densities. (**B**)The IS–PF+ strain with three different initial seeding densities. Threshold $L_{th} = 0.1 * L_{max}$. Shaded region corresponds to one standard deviation of three technical replicates. Both axes are set to a logarithmic scale.

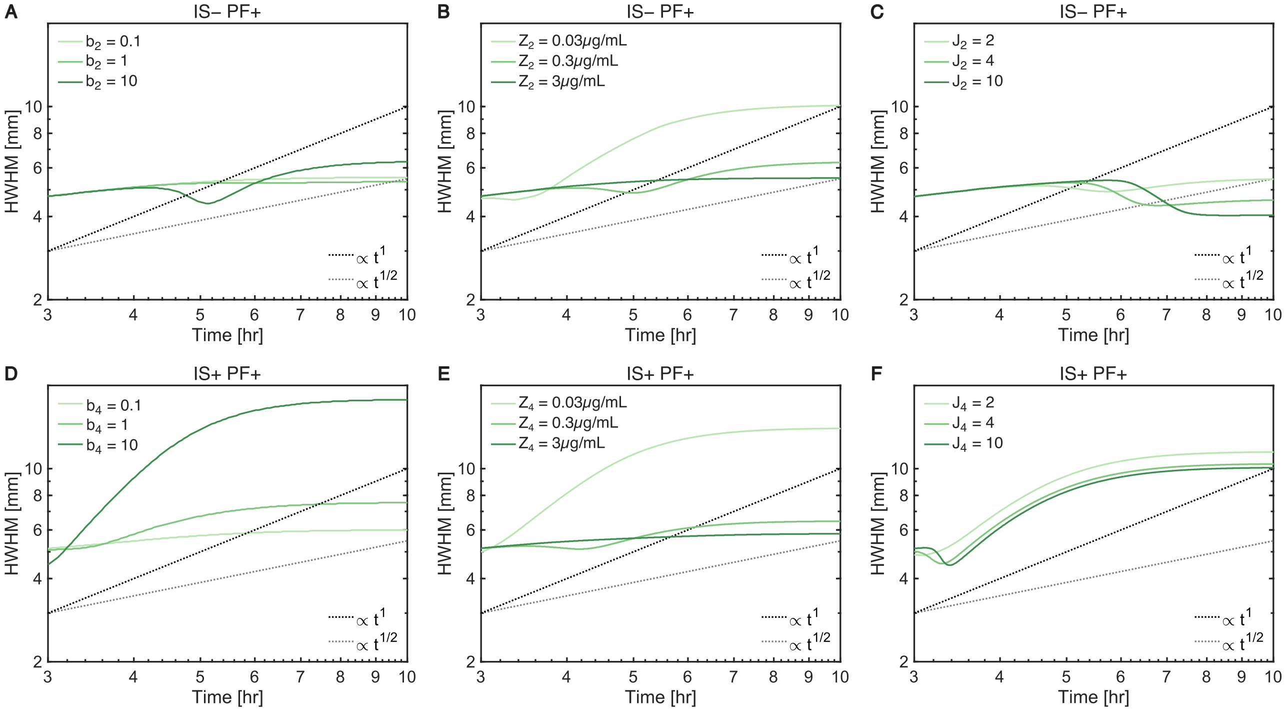


Figure S9: Parameter calibration of mathematical model. (**A-C**): Effect of modulating the free parameters in the IS–PF+ strain. (**D-F**): Effect of modulating the free parameters in the IS+PF+ strain.

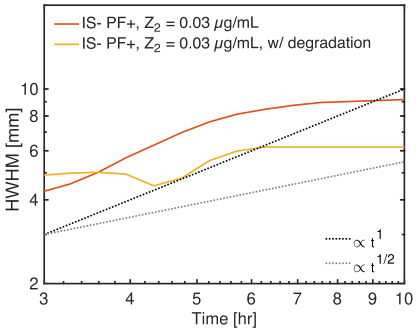


Figure S10: Wavefront dynamics of positive feedback circuit with AraC degradation, Equation 17.

### Linear analysis of the mathematical model

Here we present a linear analysis of our mathematical model to elucidate the propagation scalings of the four circuits, as well as the effect of cell growth. For simplicity, we reduce the model to one spatial dimension, $x$ (we find numerically that whether the model is in one or two dimensions does not affect the scalings), and we neglect the $y$ dynamics (because they propagate linearly from $z$ and thus will not affect the scalings).

The linear dynamics of $s$ and $z$ in one dimension read

$$\dot{s} = Ds'', \tag{20}$$
$$\dot{z} = \alpha s + \beta z + Ez'', \tag{21}$$

where we have linearized by taking the Hill coefficients to $H_i = J_i = 1$ and assuming $s$ and $z$ are much smaller than their half-maximal values for the activation and feedback functions, allowing us to define $\alpha \equiv a_i/s_i$ and $\beta \equiv a_i b_i/z_i$. We have also neglected the cell growth function, $f(t)$. We will first analyze the scalings without this function, and then subsequently investigate its effect on the propagation.

We initialize this simplified system with no readout and with the source localized to the origin,

$$s(x,0) = \phi\delta(x), \tag{22}$$
$$z(x,0) = 0, \tag{23}$$

where $\phi$ is a function of $s_0$ and $R_0$ and sets the initial amount of the source.

The dynamics can be found by Fourier transforming in space according to

$$\hat{s}(k,t) = \int_{-\infty}^{\infty} dx\, s(x,t)e^{ikx}, \tag{24}$$
$$s(x,t) = \int_{-\infty}^{\infty} \frac{dk}{2\pi}\hat{s}(k,t)e^{-ikx}, \tag{25}$$

and similarly for $z$. Eqs. 20-23 become

$$\dot{\hat{s}} = -Dk^2\hat{s}, \tag{26}$$
$$\dot{\hat{z}} = \alpha\hat{s} + (\beta - Ek^2)\hat{z}, \tag{27}$$
$$\hat{s}(x,0) = \phi, \tag{28}$$
$$\hat{z}(x,0) = 0. \tag{29}$$

The solutions to Eqs. 26-29 are found by direct integration, using an integrating factor for $\hat{z}$,

$$\hat{s}(x,t) = \phi e^{-Dk^2 t}, \tag{30}$$
$$\hat{z}(x,t) = \frac{\alpha\phi[e^{-Dk^2 t} - e^{(\beta - Ek^2)t}]}{(E-D)k^2 - \beta}. \tag{31}$$

The back-transform of $\hat{s}$ is found by completing the square and evaluating the Gaussian integral,

$$s(x,t) = \phi\int_{-\infty}^{\infty}\frac{dk}{2\pi}e^{-Dk^2 t - ikx} = \phi\frac{e^{-x^2/4Dt}}{\sqrt{4\pi Dt}}. \tag{32}$$

The source is a spreading Gaussian, as expected. $z(x,t)$ will be evaluated individually for the four circuits next.

#### IS–PF– case

For $\beta = E = 0$, we insert Eq. 32 into Eq. 21 and integrate. The result is

$$z(x,t) = \alpha\phi\sqrt{\frac{t}{D}}\left(\frac{e^{-\zeta^2}}{\sqrt{\pi}} - \zeta\,\mathrm{erfc}\,\zeta\right), \tag{33}$$

where $\zeta \equiv x/\sqrt{4Dt}$, and $\mathrm{erfc}\,\zeta = 1 - \mathrm{erf}\,\zeta$ is the complementary error function. Locations well within the bulk of the source profile satisfy $x \ll \sqrt{Dt}$, for which $\zeta \ll 1$. To first order in $\zeta$, Eq. 33 becomes

$$z(x,t) = \alpha\phi\sqrt{\frac{t}{D}}\left(\frac{1}{\sqrt{\pi}} - \zeta\right). \tag{34}$$

Because our model is linear, the growth of $z$ is unbounded. Therefore, here and throughout, rather than defining its characteristic lengthscale using its maximum or by spatially integrating its profile, we use a fixed threshold. Specifically, we define the lengthscale as the location $x_*$ at which $z$ equals a threshold $z_*$. Applying the condition $z(x_*, t) = z_*$ to Eq. 34 and solving for $x_*$ obtains

$$x_* = \sqrt{\frac{4Dt}{\pi}}\left(1 - \frac{z_*}{\alpha\phi}\sqrt{\frac{\pi D}{t}}\right). \tag{35}$$

The second term vanishes with time, resulting in

$$x_* \to \sqrt{4Dt/\pi}. \tag{36}$$

We see that the readout spreads diffusively as $t^{1/2}$. This makes sense because it is responding at every location to the diffusing source.

#### IS+PF– case

For $\beta = 0$, we specialize to $E = D$ for simplicity. Eq. 31 becomes

$$\hat{z}(k,t) = \alpha\phi t e^{-Dk^2 t}. \tag{37}$$

The back-transform is found by completing the square and evaluating the Gaussian integral as in Eq. 32,

$$z(x,t) = \alpha\phi t\frac{e^{-x^2/4Dt}}{\sqrt{4\pi Dt}}. \tag{38}$$

The readout is a spreading Gaussian whose area increases linearly in time. Applying $z(x_*, t) = z_*$ and solving for $x_*$ obtains

$$x_* = \left[4Dt\ln\left(\frac{\alpha\phi}{2z_*}\sqrt{\frac{t}{\pi D}}\right)\right]^{1/2}. \tag{39}$$

Ignoring the log correction, which will be small, the result becomes

$$x_* \to \sqrt{4Dt}. \tag{40}$$

We see that the readout spreads as $t^{1/2}$. This makes sense because the fact that the readout itself diffuses does not significantly change the fact that, as in the previous case, it scales diffusively in response to the diffusing source.

#### IS–PF+ case

For $E = 0$, we insert Eq. 32 into Eq. 21 and integrate. The result is

$$\bar{z}(\chi,\tau) = \frac{e^{\tau-\chi}}{4}\left[\operatorname{erf}\left(\frac{2\tau-\chi}{2\sqrt{\tau}}\right)+1\right] + \frac{e^{\tau+\chi}}{4}\left[\operatorname{erf}\left(\frac{2\tau+\chi}{2\sqrt{\tau}}\right)-1\right], \tag{41}$$

where, in terms of the characteristic length $\lambda \equiv \sqrt{D/\beta}$, we have defined the dimensionless concentration $\bar{z} \equiv z/(\alpha\phi/\beta\lambda)$, length $\chi \equiv x/\lambda$, and time $\tau \equiv \beta t$.

At long times $\tau \gg 1$, we consider locations $\chi$ very close to $\tau$. We will justify post hoc that this regime is the relevant one for the profile lengthscale $\chi_*$. Defining $\epsilon \equiv \tau - \chi \ll 1$, Eq. 41 becomes

$$\bar{z}(\chi,\tau) = \frac{e^{\epsilon}}{4}\left[\operatorname{erf}\left(\frac{\tau+\epsilon}{2\sqrt{\tau}}\right)+1\right] + \frac{e^{2\tau-\epsilon}}{4}\left[\operatorname{erf}\left(\frac{3\tau-\epsilon}{2\sqrt{\tau}}\right)-1\right]. \tag{42}$$

For $\tau \gg 1$ we use the large-argument expansion $\operatorname{erf}\zeta \approx 1 - e^{-\zeta^2}/\zeta\sqrt{\pi}$ to obtain, to first order in $\epsilon$,

$$\bar{z}(\chi,\tau) = \frac{e^{\epsilon}}{2} - \frac{1}{4\sqrt{\pi}}\left(\frac{2}{\sqrt{\tau}}-\epsilon\right)e^{-\tau/4-\epsilon\sqrt{\tau}+\epsilon} - \frac{1}{4\sqrt{\pi}}\left(\frac{2}{3\sqrt{\tau}}+\epsilon\right)e^{-\tau/4+3\epsilon\sqrt{\tau}-\epsilon}. \tag{43}$$

For $\tau \gg 1$ the second and third terms vanish, leaving

$$\bar{z}(\chi,\tau) = \frac{e^{\tau-\chi}}{2}. \tag{44}$$

Applying $\bar{z}(x_*, t) = \bar{z}_*$ and solving for $\chi_*$ obtains

$$\chi_* = \tau - \ln(2\bar{r}_*), \tag{45}$$

and we see that focusing on $\chi$ values near $\tau$ is justified so long as $\bar{z}_*$ is order one. In terms of the original variables, Eq. 45 reads

$$x_* = \sqrt{\beta D}t - \sqrt{\frac{D}{\beta}}\ln\left(\frac{2z_*\sqrt{\beta D}}{\alpha\phi}\right), \tag{46}$$

We see that the readout has a linear term and a constant term. Numerically, we find that the linear term in Eq. 46 becomes dominant only at very long times. Therefore, while a ballistic scaling appears possible in this case, it is not likely to manifest at early times. Moreover, it is dependent on the feedback not saturating, as assumed upon linearizing, which is not realistic experimentally. This is likely why a ballistic scaling is not seen in the experiments.

#### IS+PF+ case

Again we specialize to $E = D$ for simplicity. Eq. 31 becomes

$$\hat{z}(k,t) = \frac{\alpha\phi}{\beta}(e^{\beta t}-1)e^{-Dk^2 t}. \tag{47}$$

The back-transform is found by completing the square and evaluating the Gaussian integral as in Eq. 32,

$$z(x,t) = \frac{\alpha\phi}{\beta}(e^{\beta t}-1)\frac{e^{-x^2/4Dt}}{\sqrt{4\pi Dt}}. \tag{48}$$

The readout is a spreading Gaussian whose area increases exponentially in time. Crucially, we will see that this exponential increase changes the scaling of the spread. Applying $z(x_*, t) = z_*$ and solving for $x_*$ obtains

$$x_* = \left\{ 4Dt \ln \left[ \frac{\alpha\phi(e^{\beta t} - 1)}{\beta z_* \sqrt{4\pi D t}} \right] \right\}^{1/2} \tag{49}$$

$$\rightarrow \left\{ 4Dt \left[ \beta t + \ln \left( \frac{\alpha\phi}{\beta z_* \sqrt{4\pi D t}} \right) \right] \right\}^{1/2}, \tag{50}$$

where the second step neglects the $1$ compared to the exponential term. Then, the second term vanishes with time, resulting in

$$x_* \rightarrow 2\sqrt{D\beta} t. \tag{51}$$

We see that the readout spreads ballistically as $t^1$. This makes sense because it is amplifying itself and diffusing—the two ingredients needed for a trigger wave. The source provides the initial impulse but would not need to diffuse for the readout to travel ballistically.

#### Cell growth

Reintroducing cell growth, we have

$$\dot{s} = Ds'', \tag{52}$$

$$\dot{z} = f(t)(\alpha s + \beta z) + Ez''. \tag{53}$$

The general solution is

$$\hat{z}(k, t) = \alpha\phi F(t) e^{-Dk^2 t} \tag{54}$$

$$\Rightarrow \quad z(x, t) = \alpha\phi F(t) \frac{e^{-x^2/4Dt}}{\sqrt{4\pi D t}}, \tag{55}$$

where

$$F(t) = e^{\beta \int_0^t dt' \, f(t')} \int_0^t dt' \, f(t') e^{-\beta \int_0^{t'} dt'' \, f(t'')}. \tag{56}$$

Applying $z(x_*, t) = z_*$ and solving for $x_*$ obtains

$$x_* = \left\{ 4Dt \left[ \ln \beta F(t) + \ln \frac{\alpha\phi}{\beta z_* \sqrt{4\pi D t}} \right] \right\}^{1/2}. \tag{57}$$

The second term vanishes with time, leaving

$$x_* \rightarrow \sqrt{4Dt \ln \beta F(t)}. \tag{58}$$

Can the exponential growth phase of $f$ cause hyperballistic spreading? To investigate this question within our linear model, we expand the exponential growth term $e^{gt}$ to linear order, giving $f(t) = 1 + gt$ for growth rate $g$. The dominant part of $F(t)$ comes from the first exponential in Eq. 56, $F(t) \sim e^{\beta t(1+gt/2)}$. Eq. 58 becomes

$$x_* \approx \sqrt{4\beta D} t \sqrt{1 + gt/2}. \tag{59}$$

We see that the readout spreads hyperballistically, with a scaling that, at long times, gets as large as $t^{3/2}$. This shows that the growth dynamics couple to the propagation dynamics to enhance the scaling, and it suggests that cell growth can indeed cause hyperballistic spread of a trigger wave.

Table S1: List of all parameters used to generate the simulation Figures 3, 4, and 5 in the main text.

| Parameter name | Value | Units | Explanation |
|---|---|---|---|
| $R$ | 30 | $\mathrm{mm}$ | Domain Size |
| $T$ | 10 | $\mathrm{hours}$ | Total Time |
| $R_0$ | 0.5 | $\mathrm{mm}$ | Initial eryC droplet size |
| $s_0$ | 2000 | $\mu\mathrm{g/mL}$ | Initial eryC concentration |
| $D$ | 500 | $\mu\mathrm{m}^2/\mathrm{s}$ | Diffusion constant, $s$ |
| $E$ | 800 | $\mu\mathrm{m}^2/\mathrm{s}$ | Diffusion constant, $z$ |
| $a_1$ | 1 | $\mu\mathrm{g/mL/s}$ | Activation strength of eryC |
| $a_3$ | 1 | $\mu\mathrm{g/mL/s}$ | Activation strength of eryC |
| $s_1$ | 8.7 | $\mu\mathrm{g/mL}$ | Half-maximal concentration of eryC |
| $s_3$ | 6.3 | $\mu\mathrm{g/mL}$ | Half-maximal concentration of eryC |
| $H_1$ | 1.6 | - | Hill coefficient for eryC |
| $H_3$ | 1.9 | - | Hill coefficient for eryC |
| $b_2$ | 3.5 | - | Feedback strength of araC |
| $b_4$ | 3.5 | - | Feedback strength of rhlI |
| $J_2$ | 2 | - | Hill coefficient for araC |
| $J_4$ | 2 | - | Hill coefficient for rhlI |
| $z_2$ | 0.5 | $\mu\mathrm{g/mL}$ | Half-maximal concentration of araC |
| $z_4$ | 0.05 | $\mu\mathrm{g/mL}$ | Half-maximal concentration of rhlI |

Table S2: Growth parameter values for different strains and optical densities (OD = 0.2, 0.6, 1.8).

| Strain | Parameter | OD = 0.2 | OD = 0.6 | OD = 1.8 |
|---|---|---|---|---|
| | $\alpha(\mathrm{h}^{-1})$ | 4 | 4 | 4 |
| IS–PF– | $g\,(\mathrm{h}^{-1})$ | 1.58 | 1.53 | 2.18 |
| | $n_0$ (log) | 4.05 | 4.98 | 5.71 |
| | $C$ (log) | 9.87 | 10.57 | 10.08 |
| | $\lambda\,(\mathrm{h})$ | 0.97 | 0.62 | 0.83 |
| IS–PF+ | $g\,(\mathrm{h}^{-1})$ | 1.40 | 1.52 | 1.87 |
| | $n_0$ (log) | 3.70 | 5.39 | 5.85 |
| | $C$ (log) | 10.77 | 10.16 | 10.09 |
| | $\lambda\,(\mathrm{h})$ | 0.98 | 1.44 | 1.29 |
| IS+PF– | $g\,(\mathrm{h}^{-1})$ | 1.86 | 2.36 | 2.06 |
| | $n_0$ (log) | 4.15 | 4.70 | 5.74 |
| | $C$ (log) | 10.68 | 10.67 | 11.08 |
| | $\lambda\,(\mathrm{h})$ | 0.99 | 1.08 | 0.71 |
| IS+PF+ | $g\,(\mathrm{h}^{-1})$ | 1.96 | 1.91 | 2.22 |
| | $n_0$ (log) | 5.22 | 5.46 | 6.54 |
| | $C$ (log) | 10.31 | 10.64 | 10.67 |
| | $\lambda\,(\mathrm{h})$ | 1.51 | 0.99 | 1.13 |

Table S3: Plasmids used in this study. Ori: origin of replication.

| **Plasmids** | **Genes** | **Ori** | **Resistance** |
|---|---|---|---|
| pMDW476 | pmph-BCD7-rhlI-ssrA prhl-BCD10-sfyfp laclq-mphR-eryR | pSC101 | Spc |
| pMDW621 | Prhl HHRZ-supP | p15A | Kan |
| pMDW677 | Prhl-BCD2-rhlI.2TAG-ssrA HHRZ-supP | p15A | Kan |
| pMDW502 | pmph-BCD7-araC-ssrA pBAD-BCD10-sfyfp laclq-mphR-eryR | pSC101 | Spc |
| pMDW628 | PBAD HHRZ-supP | p15A | Kan |
| pMDW678 | PBAD-BCD2-araC.2TAG-ssrA HHRZ-supP | p15A | Kan |
| pMDW146 | prhl BCD10 rhlI-ssrA | p15A | Amp |
| pMDW503 | pBAD BCD10 arac-ssrA | p15A | Kan |
